# A Novel CREBBP/p300 Activator, YF2, Enhances Cytotoxic and Immune-Mediated Responses in B-Cell Lymphoma

**DOI:** 10.1101/2025.03.13.642871

**Authors:** Ted B. Piorczynski, Yuxuan Liu, Brian Estrella, Edd C. Ricker, Manuel Pazos, Yulissa Gonzalez, Seda S. Tolu, Yun Kyoung Ryu Tiger, Aleksandar Obradovic, Chang Liu, Charles Karan, Serge Cremers, Renu Nandakumar, Barry Honig, Howook Hwang, Neil L. Kelleher, Jeannie M. Camarillo, Nebiyu Ali Abshiru, Jennifer Effie Amengual

## Abstract

Germinal center B-cell lymphomas frequently exhibit monoallelic loss-of-function mutations in the histone acetyltransferases CREBBP and p300, which contribute to lymphoma development, disrupt normal germinal center biology, and promote immune evasion. This study evaluates YF2, a histone acetyltransferase activator that increases CREBBP/p300 activity, as a potential therapeutic approach for B-cell lymphoma. YF2 binds the bromodomain of CREBBP/p300, increasing their auto-acetylation and enzymatic activity. It also induces cytotoxicity in B-cell lymphoma cell lines, with a stronger response in those carrying *CREBBP*/*EP300* mutations. Additionally, YF2 increases the acetylation of key CREBBP/p300 substrates, including H3K27, p53, and BCL6, leading to enhanced apoptosis and altered B-cell differentiation. YF2 is well tolerated *in vivo* and extends survival in both cell line- and patient-derived xenograft mouse models. Moreover, YF2 exposure upregulates antigen-presentation markers and reshapes the immune microenvironment, strengthening responses to immune checkpoint blockade. Taken together, these findings support pharmacologic activation of CREBBP/p300 with YF2 as a compelling therapeutic strategy for B-cell lymphoma.

**Significance:** YF2 stimulates CREBBP/p300 activity to normalize B-cell differentiation and enhance immune recognition, offering a potential approach to limit lymphoma progression and strengthen immunotherapy responses.

## Introduction

Epigenetic dysregulation is a key driver of lymphoma pathogenesis^1–3^. Diffuse large B-cell lymphoma (DLBCL) and follicular lymphoma (FL), the most common subtypes, originate from germinal center (GC) B cells that acquire oncogenic epigenetic mutations^4,5^. Inactivating mutations in the histone acetyltransferases (HATs) CREBBP and its paralogue p300 are common in GC-derived lymphomas, affecting approximately 40% of DLBCL^4,6,7^ and 60% of FL cases^8,9^. CREBBP and p300 are broad transcriptional coactivators with extensive protein interactions and substrates^10^, including the tumor suppressor p53^11,12^ and the proto-oncogene BCL6^13^; acetylation of p53 enhances its tumor-suppressive activity and protects it from degradation, whereas acetylation of BCL6 neutralizes its oncogenic function^14^. CREBBP and p300 also regulate GC dynamics and major histocompatibility complex (MHC)-associated antigen presentation^15^. Loss of CREBBP and/or p300 disrupts these processes, driving lymphomagenesis, immune evasion, and disease progression^16–18^.

Mutations in *CREBBP* and *EP300* are typically monoallelic, resulting in haploinsufficiency despite retention of the wild-type (WT) allele^8,19^. This functional allele, together with a potentially intact paralogue, provides an opportunity to leverage the unaltered HAT protein to overcome the pathogenic impact of the mutated enzyme. Epigenetic therapies such as histone deacetylase inhibitors (HDACi) and hypomethylating agents have emerged as potential therapeutic approaches to counteract lymphoma proliferation^20,21^, while HAT bromodomain inhibitors exploit synthetic lethality^22,23^. Phase 1 clinical trials of bromodomain inhibitors have been conducted in various diseases^24,25^, but they have not yet been evaluated in lymphomas or other malignancies harboring HAT mutations. To date, few therapeutic strategies have aimed to *activate* the remaining functional HAT allele or the unaffected paralogue to correct the epigenetic dysregulation characteristic of *CREBBP*/*EP300*-mutated lymphomas.

YF2 is a commercially available HAT activator originally synthesized by the Columbia University Organic Chemistry Collaborative Center^26^. The present study evaluates YF2 as a therapeutic strategy for GC B-cell (GCB) lymphomas, particularly those harboring *CREBBP*/*EP300* mutations. YF2 induces cytotoxicity preferentially in *CREBBP*/*EP300*-mutated lymphoma cell lines, promotes B-cell differentiation, modulates the immune microenvironment, and demonstrates efficacy across multiple lymphoma mouse models, supporting HAT activation as a therapeutic strategy in epigenetically dysregulated malignancies.

## Methods

### Ethical approval

All animal procedures were approved by the Columbia University Institutional Animal Care and Use Committee. Human samples were obtained following written informed consent under protocols approved by the Columbia University Institutional Review Board.

### Drug synthesis and acquisition

CTPB was purchased from MedChemExpress (#A18558). YF2, JF1, JF10, JF16, and RP102 were initially synthesized at the Organic Chemistry Collaborative Center at Columbia University; subsequent experiments used commercially sourced YF2 (MedChemExpress #HY-16531A), which exhibited equivalent potency and reactivity to the synthetic compound.

### Cell culture

Cell line mutations were verified through prior studies^7,19^ and DepMap^27^ (https://depmap.org/portal). A20 (RRID: CVCL_1940), HBL-1 (RRID: CVCL_4213), OCI-Ly1 (RRID: CVCL_1879), PfieTer (RRID: CVCL_3326), RIVA (RRID: CVCL_1885), SU-DHL-2 (RRID: CVCL_9550), SU-DHL-6 (RRID: CVCL_2206), and Toledo (RRID: CVCL_3611) were obtained from American Type Culture Collection (ATCC); OCI-Ly7 (RRID: CVCL_1881), OCI-Ly10 (RRID: CVCL_8795), and SU-DHL-10 (RRID: CVCL_1889) from Leibniz Institute DSMZ; BJAB (RRID: CVCL_5711), SU-DHL-4 (RRID: CVCL_0539), and U-2932 (RRID: CVCL_1896) from the laboratory of Laura Pasqualucci; and DB (RRID: CVCL_1168) and Karpas-422 (RRID: CVCL_1325) from the laboratory of Chao Lu. A20 cells underwent whole-exome sequencing and SNP/indel analysis by GENEWIZ (Azenta Life Sciences). Cells were cultured in RPMI 1640 (Thermo Fisher Scientific #11875093) or IMDM (Gibco #12440053) supplemented with 10–20% fetal bovine serum (FBS; Gibco #A5670801) as follows: RPMI (10% FBS): BJAB, DB, HBL-1, Karpas-422, Pfeiffer, RIVA, SU-DHL-4, SU-DHL-6, SU-DHL-10, Toledo, U-2932; RPMI (10% FBS + 50 µM 2-mercaptoethanol): A20; RPMI (20% FBS): Farage, SU-DHL-2; IMDM (10% FBS): OCI-Ly7; IMDM (20% FBS): OCI-Ly1, OCI-Ly10.

Cells were cultured in RPMI 1640 (Thermo Fisher Scientific #11875093) or IMDM (Gibco #12440053) supplemented with 10–20% fetal bovine serum (FBS; Gibco #A5670801) under standard conditions (37 °C, 5% CO₂), with media formulations tailored according to provider recommendations for each cell line. Cells were passaged every 2–3 days, authenticated by ATCC, and routinely tested for mycoplasma contamination.

### Cell viability assay

Cells were plated at 50 cells/µL in 96-well plates and treated with CTPB, YF2, and/or A-485 (MedChemExpress #HY-107455) diluted in dimethyl sulfoxide (DMSO; Thermo Fisher Scientific #85190). Viability was assessed at indicated time points using Trypan blue exclusion or CellTiter-Glo 2.0 (Promega #G924) following 10 min incubation at room temperature. Luminescence was measured using a GloMax Explorer plate reader (Promega). The half-maximal inhibitory concentration (IC_50_) values were calculated using the drc package^28^ (v3.0-1) in R (v4.3.0), and data represent the mean of three technical replicates unless otherwise stated.

### Surface plasmon resonance (SPR)

SPR was performed by Ichor Life Sciences using an NTA sensor chip. p300 bromodomain protein (Active Motif #31372) was immobilized (∼2032 RU) following Ni²⁺ recharging and amine coupling and blocked with ethanolamine. Binding was assessed by single-cycle kinetics at 25 °C using twofold serial dilutions. Association and dissociation times were 120 s and 1200 s for Inobrodib and YF2, respectively, and data were processed by reference subtraction, buffer blanking, and solvent correction (0.4–1.6% DMSO).

### Medium-throughput compound screening

Cytotoxicity of 31 epigenetic compounds was assessed in SU-DHL-10, SU-DHL-2, Farage, and HBL-1 cells. Romidepsin was purchased from MedChemExpress (#HY-15149), RA series compounds were provided by Appia Pharmaceuticals, and all other compounds were provided by the Organic Chemistry Collaborative Center at Columbia University. Cells were plated in triplicate at 50 cells/µL using a D300 Digital Dispenser (HP), treated with compounds, and viability was measured on days 3 and 6 using CellTiter-Glo 2.0. IC₅₀ values were calculated as described above.

### Ligand-protein binding prediction

Predicted binding of YF2 to p300 was performed using the LBias algorithm^29^. This method identifies structurally similar proteins, aligns ligand-bound templates, and scores interaction compatibility using the p300 crystal structure from the Protein Data Bank^30^.

### Cell-free thermal shift assay

Thermal stability of p300 was evaluated using the Protein Thermal Shift Kit (Thermo Fisher Scientific #4462263). Recombinant p300 (Sigma-Aldrich #03188) protein covering the HAT domain and p300 (Active Motif #31372) protein encompassing the bromodomain (both at 1 µM) were incubated with 0–50 µM YF2 at room temperature for 10 min. Select samples were incubated with 10 µM Inobrodib (MedChemExpress #HY-111784) as a positive control. Samples were combined with assay buffer and dye, heated from 25–99 °C (ramp rate of 0.05 °C/s) using a QuantStudio 3 real-time PCR system (Applied Biosystems), and fluorescence was monitored to determine melting temperatures (Tₘ) using Protein Thermal Shift Software (v1.4; Applied Biosystems).

### HAT auto-acetylation

CREBBP/p300 proteins (1 µM; Active Motif #31990 and #81893, respectively) were deacetylated using 100 ng SIRT2 (Active Motif #31528) and 5 mM NAD⁺ at 30 °C for 3 h, followed by quenching with nicotinamide. Auto-acetylation reactions were performed with acetyl-CoA and incubated with 50 µM YF2 or vehicle at 30 °C for 1–10 min. Reactions were terminated with loading buffer at defined time points and analyzed by immunoblotting as described below.

### Cell-free acetylation

Recombinant CREBBP or p300 (100 nM; Active Motif #31590 and #31124, respectively) was incubated with 0.1–100 µM YF2 for 30 min at 30 °C, followed by the addition of acetyl-CoA and substrates: histone H3 (1 µg; New England Biolabs #M2507S), p53 (100 ng; Sigma-Aldrich #14-865), or BCL6 (100 ng; MyBioSource #MBS7135965). Reactions were incubated for 1 h at 30 °C, terminated, and analyzed by immunoblotting. EC₅₀ values were calculated using the drc package^28^ (v3.0-1) in R (v4.3.0).

### Kinase screening

Kinase profiling was performed by DiscoverX using 10 µM YF2. Binding interactions were quantified using DNA-tagged kinases and qPCR-based detection following affinity selection.

### HAT activity assay

HAT activity was measured using a commercial kit (Active Motif #56100). Recombinant CREBBP (Active Motif #31590), p300 (Active Motif #81093), KAT2A (Active Motif #31591), and KAT2B (Active Motif #81142) proteins were incubated with the kit-supplied histone peptides and 0–50 µM YF2 for 30 min at room temperature. Fluorescence was measured after YF2 treatment on a GloMax Explorer plate reader (Promega), and activity was quantified relative to controls.

### Primary sample collection and treatment

Peripheral blood mononuclear cells (PBMCs) were isolated by Ficoll density centrifugation (GE Healthcare #17144002) using SepMate tubes (STEMCELL Technologies #85415). Cells were washed, red blood cells lysed using ACK Lysing Buffer (Gibco #A10492-01), and remaining PBMCs cultured in RPMI + 20% FBS. Cells were treated with 0–10 µM YF2 or vehicle for 6 days before viability assessment.

### CRISPR–Cas9 knockout generation

sgRNAs targeting CREBBP or p300 were purchased from Synthego, complexed with Cas9, and nucleofected into OCI-Ly7 cells using the 4D-Nucleofector System (Lonza Bioscience) using the CM-137 electroporation program with Solution F. Single-cell clones were isolated by limiting dilution and expanded for validation and testing.

### Mass spectrometry

SU-DHL-6 cells were treated *in vitro* with 6 µM YF2 or vehicle for 48 h, while *in vivo* samples were obtained from SU-DHL-6 xenograft tumors following 5 days of YF2 treatment (40 mg/kg). Cells and tumors were washed, flash frozen, and stored at –80 °C before processing. Samples were resuspended in 0.2 N sulfuric acid and incubated at room temperature for 1 h, followed by centrifugation to remove debris. Histones were precipitated with 20% (v/v) trichloroacetic acid on ice, pelleted, and washed sequentially with acidified acetone and 100% acetone before drying and storage at –20 °C. Histones were propionylated and digested as described previously^31^, with a single round of propionylation performed before and after digestion. Targeted LC–MS/MS was performed on a TSQ Quantiva Triple-Stage Quadrupole Mass Spectrometer (Thermo Fisher Scientific), and data were analyzed using Skyline^32^. All proteomics experiments were conducted by the Northwestern Proteomics Core Facility.

### Histone extraction

Cells were treated with YF2 or vehicle for three days, lysed in TEB buffer (PBS containing 2% protease inhibitor cocktail, 0.5% Triton X-100, and 0.02% sodium azide) at 4°C for 10 min, and nuclei were isolated. Histones were acid-extracted using 0.2 N hydrochloric acid at 4°C overnight. The next day, histones were precipitated with 70% perchloric acid at 4°C for 1 h, washed with acetone, and stored at −80 °C for downstream analysis.

### Immunoblotting

Histone extract and whole-cell lysate concentrations were measured using Pierce BCA Protein Assay Kits (Thermo Fisher Scientific #23225). Equal amounts of protein were mixed with sample buffer containing 5% (v/v) 2-mercaptoethanol, boiled at 95°C for 5 min, loaded on Mini-Protean TGX 4-15% gels (Bio-Rad #4568084), and run using a Mini-Protean Tetra Cell System (Bio-Rad #1658004). Proteins were transferred onto PVDF membranes (Cytiva #10600021) using the Trans-Blot Turbo Transfer System (Bio-Rad #1704150). Membranes were blocked in 5% bovine serum albumin (BSA) for acetylated targets or 5% milk for all others at room temperature for 1 h, followed by overnight incubation with primary antibodies (Table 1) diluted in TBST (tris-buffered saline containing 0.1% Tween 20) at 4°C. After primary antibody incubation, membranes were incubated with HRP-linked anti-rabbit (Cell Signaling Technology #7074; RRID: AB_2099233) or anti-mouse (Cell Signaling Technology #7076; RRID: AB_330924) secondary antibodies for 1 h at room temperature. Blots were developed using SuperSignal West Pico Plus substrate (Thermo Fisher Scientific #34580) and imaged on film, and densitometry was performed using ImageJ software (v1.54).

**Table 1:** Primary antibodies used for immunoblotting.

| Target | Vendor | Catalog number | RRID |
| --- | --- | --- | --- |
| CREBBP | Cell Signaling Technology | 7389 | AB_2616020 |
| p300 | Cell Signaling Technology | 54062 | AB_2799450 |
| Acetyl-CREBBP/p300 | Cell Signaling Technology | 4771 | AB_2262406 |
| H3K18ac | Cell Signaling Technology | 9675 | AB_331550 |
| H3K27ac | Cell Signaling Technology | 4353 | AB_10545273 |
| H3 | Cell Signaling Technology | 4499 | AB_10544537 |
| Acetyl-lysine | Cell Signaling Technology | 9441 | AB_331805 |
| Acetyl-p53 | Cell Signaling Technology | 2525 | AB_330083 |
| p53 | Cell Signaling Technology | 2527 | AB_10695803 |
| p21 | Cell Signaling Technology | 2947 | AB_823586 |
| PUMA | Cell Signaling Technology | 12450 | AB_2797920 |
| BCL6 | Cell Signaling Technology | 4242 | AB_2063431 |
| PRDM1 | Cell Signaling Technology | 9115 | AB_2169699 |
| Mouse MHC-I | Invitrogen | PA5-115363 | AB_2899999 |
| Human MHC-I | Invitrogen | MA5-35712 | AB_2849612 |
| Mouse MHC-II | FabGenniX | MHCII-101AP | Not available |
| Human MHC-II | Cell Signaling Technology | 68258 | Not available |
| B2M | Abcam | AB75853 | AB_1523204 |
| GAPDH | Cell Signaling Technology | 5174 | AB_10622025 |

### Generation of histone deacetylase inhibitor (HDACi)-resistant cell lines

OCI-Ly7 and SU-DHL-6 cells were treated with increasing concentrations of romidepsin for 6 days, after which IC₁₀ and IC₅₀ values were determined. Cells were continuously cultured at IC₁₀ doses for 6 days with periodic recalculation until a ∼10-fold shift in IC₅₀ was achieved.

### Pharmacokinetics

SCID/beige mice (6–8 weeks old; Charles River Laboratories; RRID: IMSR_CRL:250) were subcutaneously engrafted with 1 × 10⁷ SU-DHL-6 cells in 50% Matrigel. Tumors were measured three times weekly using calipers and volumes calculated as (length × width²)/2. Mice were enrolled once their tumors reached a volume of 80 mm³, then treated with YF2 (40 or 60 mg/kg) via intraperitoneal (i.p.) injection once daily for 5 days. Tumors were homogenized in LC–MS-grade water at 4 °C and processed alongside plasma samples to compare systemic and intratumoral drug exposure. YF2 was extracted using a 3:1 (v/v) acetonitrile:methanol solution, followed by centrifugation and LC–MS/MS analysis on an Agilent UHPLC–triple quadrupole platform. Calibration curves (1 ng/mL to 1 µg/mL) were generated in control serum and processed identically to samples. Pharmacokinetic parameters were calculated using Phoenix WinNonlin (v8.5; Certara).

### Xenogeneic lymphoma mouse models

SCID/beige mice (6–8 weeks old) were subcutaneously engrafted with either 1 × 10⁷ SU-DHL-6 cells or TM00368 patient-derived DLBCL cells in 50% Matrigel to generate cell line–derived xenograft and PDX models, respectively. Once tumors reached 80 mm³, mice were randomized to receive vehicle (saline) or YF2 (40 mg/kg, i.p., 5 days/week). Treatment duration was 4 weeks for SU-DHL-6 xenografts and 8 weeks for PDX models to account for differing growth kinetics. Mice were euthanized if body weight decreased by >20% or tumor volume exceeded 2,000 mm³, in accordance with institutional guidelines.

### Gene signaling arrays

Cells were seeded at 50 cells/µL and treated with 10 µM YF2 or vehicle for 24 h. RNA was isolated, reverse transcribed, and analyzed using the TaqMan Array Human p53 Signaling Pathway array (Applied Biosystems #4414168) or the TaqMan Array Human Antigen Processing and Presentation by MHCs plate (Applied Biosystems #4414104). qPCR was performed on a QuantStudio 3 system (Applied Biosystems) under standard cycling conditions, and data from technical replicates were averaged prior to analysis using ExpressionSuite (v1.3; Thermo Fisher Scientific).

### Chromatin immunoprecipitation

SU-DHL-6 cells were treated with 10 µM YF2 or vehicle for 24 h, washed with PBS, fixed with 4% paraformaldehyde for 10 min, and snap frozen. Chromatin immunoprecipitation (ChIP) was performed using the ChIP-IT Express Enzymatic Kit (Active Motif #53035) with minor modifications. Cell pellets were subjected to hypotonic swelling and lysis, followed by Dounce homogenization to release nuclei, which were pelleted and enzymatically digested. Sheared DNA (10 µg) was incubated overnight at 4 °C with 5 µg of H3K27ac antibody (Cell Signaling Technology #4353; RRID: AB_10545273) or 5 µg of IgG control (Active Motif 101226), and the eluted DNA was purified using a commercial kit (Active Motif #58002). qPCR was performed on a QuantStudio 3 Real-Time PCR System (Applied Biosystems) under standard cycling conditions, and samples were run in triplicate and averaged before analysis. All primers were purchased from Integrated DNA Technologies: *p21* forward 5’–GAAGCATGTGACAATCAACAACTC, reverse 5’–AAGCATCTTGAGGCCAGAAT; *HLA-B* forward 5’–GGAGGGAAATGGCCTCTG, reverse 5’–GGACACGGAGGTGTAGAAATAC; *HLA-C* forward 5’–GAGACCTGGGCCTGTGA, reverse 5’–GTCGAAATACCTCATGGAGTGG.

### Bulk RNA sequencing

SU-DHL-6 and Karpas-422 cells were treated with 12 µM and 8 µM YF2, respectively, or vehicle. Cells were retreated on day 3 following a 1:1 dilution with fresh media and harvested on day 7 (SU-DHL-6) or day 6 (Karpas-422), then washed, flash frozen, and stored at –80 °C. RNA was purified using the PureLink RNA Mini Kit (Invitrogen #12183018A) with on-column DNase treatment, and RNA quality was assessed by the Agilent 2100 Bioanalyzer (Agilent Technologies), with all samples exhibiting RIN > 9. mRNA libraries were prepared from 500 ng total RNA using the TruSeq Stranded mRNA Library Prep kit (Illumina #20020595), including poly(A) selection, fragmentation, cDNA synthesis, adapter ligation, and PCR amplification with KAPA HiFi HotStart ReadyMix (Roche #KK2602). Libraries were sequenced as paired-end 75-bp reads on an Element AVITI System (Element Biosciences) at the Columbia Genome Center. Sequencing data were processed in R (v4.3.0), with quality control by FastQC^33^ (v0.11.9), alignment to the GRCh38 genome using Rsubread^34^ (v2.14.2), and differential expression analysis performed using DESeq2^35^ (v1.40.1). Genes with an adjusted *P* value < 0.05 and absolute log_2_(fold change) > 0.5 were considered differentially expressed. Gene annotation and enrichment analyses were performed using BioMart^36^ (v2.56.0), ClusterProfiler^37^ (v4.8.1), and DisGeNET^38^, and data visualization was conducted using ggplot2 (v3.4.2) and ComplexHeatmap^39^ (v2.16.0).

### Cell cycle analysis

Cells were treated with 0–20 µM YF2 for 6 days, fixed in 70% ethanol, and stained with PI/RNase solution (Invitrogen #F10797). Samples were analyzed by flow cytometry on a Cytek Aurora system (Cytek Biosciences). Cell cycle distributions were quantified using FlowJo (v10.10; BD Biosciences)

### Apoptosis assay

Cells were treated with 0–20 µM YF2 for 6 days and stained with Annexin V and 7-AAD (BioLegend #640922). Apoptotic populations were quantified by flow cytometry using a Cytek Aurora system (Cytek Biosciences). Data were analyzed using FlowJo (v10.10; BD Biosciences).

### BH3 profiling

BH3 profiling was performed as described previously^40^. Cells were treated with 10 µM YF2 for 24 h and exposed to BH3 peptides to assess mitochondrial priming. Mitochondrial outer membrane permeabilization was monitored over 90 min using fluorescence-based detection on the GloMax Explorer (Promega).

### Syngeneic lymphoma mouse model

BALB/c mice (6–8 weeks old; Taconic Biosciences; RRID: IMSR_TAC:BALB) were immunized with 1 × 10⁸ sheep red blood cells (Colorado Serum Company #31112) and treated the following day with saline or YF2 (50 mg/kg, i.p., 5 days/week) for 2 weeks, after which blood and spleens were collected. For tumor studies, mice were engrafted subcutaneously with 1 × 10⁷ A20 lymphoma cells, and tumor growth was monitored by caliper measurements. Once tumors reached 80 mm³, mice were immunized with sheep red blood cells and randomized into four groups: vehicle, YF2 (50 mg/kg, i.p., 5 days/week), anti–PD-L1 antibody (200 µg/mouse, weekly; Bio X Cell #BE0101; RRID: AB_10949073), or combination therapy. Treatment continued for 4 weeks, with a subset of mice euthanized on day 14 for flow cytometry and spatial transcriptomic analyses. Tumor volume and body weight were measured three times per week, and mice were euthanized upon >20% weight loss or tumor volume exceeding 2,000 mm³, in accordance with institutional guidelines.

### Flow cytometry

Murine blood was diluted 1:1 with blocking buffer (PBS containing 1% BSA and 2 mM EDTA). Samples were stained with antibodies for 20 min at room temperature in the dark, followed by red blood cell lysis using BD FACS Lysing Solution (BD Biosciences #349202), washing, and resuspension in blocking buffer for analysis. Splenic and tumor tissues were collected, minced on ice, and processed similarly. Tissue homogenates were lysed, stained with antibodies for 20 min at room temperature, washed, and resuspended in blocking buffer. Cell lines were washed with PBS, stained with antibodies, and resuspended in blocking buffer. Samples were analyzed on a five-laser Cytek Aurora flow cytometer (Cytek Biosciences), and data were processed using FlowJo (v10.10; BD Biosciences). All antibody concentrations were optimized prior to use (Table 2).

**Table 2:** Antibodies used for flow cytometry.

| Target | Vendor | Catalog number | RRID |
| --- | --- | --- | --- |
| Zombie NIR | BioLegend | 423105 | AB_2757961 |
| MHC-I (FITC) | BioLegend | 125508 | AB_1236470 |
| Human MHC-I (FITC) | Invitrogen | 11-9983-42 | AB_1633403 |
| MHC-II (PE/Cyanine5) | BioLegend | 107612 | AB_313327 |
| Human MHC-II (APC) | BioLegend | 361714 | AB_2750316 |
| B220 (APC/Cyanine7) | BioLegend | 103224 | AB_313007 |
| CD19 (BV650) | BioLegend | 115541 | AB_11204087 |
| CD138 (PE) | BioLegend | 142504 | AB_10916119 |
| CXCR4 (BV711) | BioLegend | 146517 | AB_2687244 |
| IgG1 (BV421) | BioLegend | 406616 | AB_2562234 |
| IgD (PerCP) | BioLegend | 405736 | AB_2563346 |
| CD21/CD35 (Alexa Fluor 700) | BioLegend | 123432 | AB_2860650 |
| CD23 (PE/Dazzle 594) | BioLegend | 101634 | AB_2687194 |
| GL7 (PerCP/Cyanine5.5) | BioLegend | 144610 | AB_2562979 |
| CD38 (PE/Cyanine7) | BioLegend | 102718 | AB_2275531 |
| CD86 (BV510) | BioLegend | 105039 | AB_2562370 |
| CD3 (PE/Cyanine5) | BioLegend | 100274 | AB_2894410 |
| CD4 (BUV805) | BD Biosciences | 741912 | AB_2871226 |
| CD8a (BV785) | BioLegend | 100750 | AB_2562610 |
| CD44 (APC/Cyanine7) | BioLegend | 103028 | AB_830785 |
| CD25 (Alexa Fluor 700) | BioLegend | 102024 | AB_493709 |
| CD274 (BV785) | BioLegend | 124331 | AB_2629659 |
| IgM (APC) | BioLegend | 406509 | AB_315059 |
| CD11b (BUV395) | BD Biosciences | 563553 | AB_2738276 |
| CXCR5 (BUV395) | BD Biosciences | 563980 | AB_2738521 |
| CD335 (BUV737) | BD Biosciences | 612805 | AB_2870131 |
| TIGIT (BV421) | BioLegend | 142111 | AB_2687311 |
| CD226 (BV510) | BioLegend | 133607 | AB_2715921 |
| B220 (BV605) | BioLegend | 103244 | AB_2563312 |
| LAG3 (BV650) | BD Biosciences | 740560 | AB_2740261 |
| VISTA (BV711) | BD Biosciences | 742724 | AB_2741000 |
| CD62L (PE) | BioLegend | 104408 | AB_313095 |
| CD45 (PerCP) | BioLegend | 103130 | AB_893339 |
| NK-1.1 (PerCP/Cyanine5.5) | BioLegend | 156526 | AB_2894655 |
| CD127 (PE-CF594) | BD Biosciences | 562419 | AB_11153131 |
| PD-1 (PE/Cyanine7) | BioLegend | 135216 | AB_10689635 |
| Tim-3 (APC) | BioLegend | 134008 | AB_2562998 |

### Immunohistochemistry

Murine spleens and tumors were fixed in 4% paraformaldehyde overnight at 4 °C, transferred to 70% ethanol, and processed into FFPE sections (5 µm) by the Columbia University Molecular Pathology Shared Resource. Sections were deparaffinized, rehydrated, and treated for endogenous peroxidase activity (Abcam #87449), followed by antigen retrieval (Abcam #AB93684). Slides were blocked (Abcam #86553) for 30 min at room temperature and incubated overnight at 4 °C with CD8 (Cell Signaling Technology #98941S; RRID: AB_2756376), BCL6 (Novus Biologicals #NBP3-07526; RRID: AB_3554916), CD86 (Cell Signaling Technology #19589; RRID: AB_2892094), or IRF-4 antibodies (Cell Signaling Technology #62834; RRID: AB_2877647). The next day, slides were sequentially incubated with HRP conjugate (Abcam #88364), DAB substrate (BD #550880), and hematoxylin. Slides were then dehydrated, cleared, mounted, and imaged using an Aperio AT2 scanner (Leica Biosystems). Images were analyzed using FlowJo (v10.10; BD Biosciences).

### Spatial transcriptomics

Murine spleens and tumors were fixed in 4% paraformaldehyde overnight at 4 °C, transferred to 70% ethanol, and processed into FFPE sections (5 µm) by the Columbia University Molecular Pathology Shared Resource. Sections were submitted to the Mount Sinai Genomics Core for library preparation and sequencing using the Visium Spatial Gene Expression for FFPE platform. Sequenced data were processed using SpaceRanger (v4.0.0; 10X Genomics), normalized with sctransform in Seurat^41^ (v5.1.0), and batch-corrected and integrated using anchor-based integration on the top 2,000 highly variable genes. Integrated datasets were reduced using the top 50 principal components, and unsupervised clustering was performed using the Louvain algorithm with resolution optimized by silhouette scoring^42^. Differential expression between clusters was assessed using MAST^43^ (v1.33.0), and cell types were annotated using canonical markers and SingleR with the ImmGen reference^44^. Spatial cluster identities were projected back onto tissue maps, and functional enrichment was performed using WebGestalt against WikiPathways gene sets^45^. Cell type abundance differences between treatment groups were assessed using Fisher’s exact test with Benjamini–Hochberg correction. Spatial deconvolution was performed using STdeconvolve^46^ (v3.20), with model parameters optimized to minimize perplexity and low-abundance cell states. Putative T cell populations were manually validated based on marker gene expression across inferred topics.

### Statistical analyses

All statistical analyses were performed in R (v4.3.0). Data were assessed for normality (Shapiro–Wilk test) and homoscedasticity (Levene’s test), and all measurements were taken from biologically independent samples. Data were plotted using ggplot2 (v3.4.2). Asterisks (*) denote statistical significance: * = *P* < 0.05, ** = *P* < 0.01, and *** = *P* < 0.001.

## Results

### YF2 induces cytotoxicity in B-cell lymphoma, exhibiting heightened e:ects in CREBBP/EP300-deficient cells

A library of 31 HAT-activating compounds was designed and synthesized using N-(4-chloro-3-trifluoromethyl-phenyl)-2-ethoxy-6-pentadecyl-benza-mide (CTPB) as a template by the Columbia University Organic Chemistry Collaborative Center^26^. The cytotoxic effects of these compounds were evaluated by medium-throughput screening across four DLBCL cell lines (Supplementary Fig. S1a). From this initial screen, five effective analogs sharing the N-phenyl benzamide scaffold were selected for further study (Supplementary Fig. S1b), and cytotoxicity was evaluated across an expanded panel of 10 B-cell lymphoma cell lines harboring various *CREBBP*/*EP300* mutations (Supplementary Fig. S1c). YF2 (Fig. 1a) was selected as the lead compound based on its selective toxicity toward *CREBBP*- and *EP300*-mutated cell lines (Supplementary Fig. S1d). To directly compare the efficacy of YF2 to its parent compound, two *CREBBP*+*EP300*-mutated GCB lymphoma cell lines were treated with increasing concentrations of YF2 or CTPB. YF2 induced greater cytotoxicity than CTPB at equivalent concentrations in both cell lines (Fig. 1b).

**Fig. 1:**
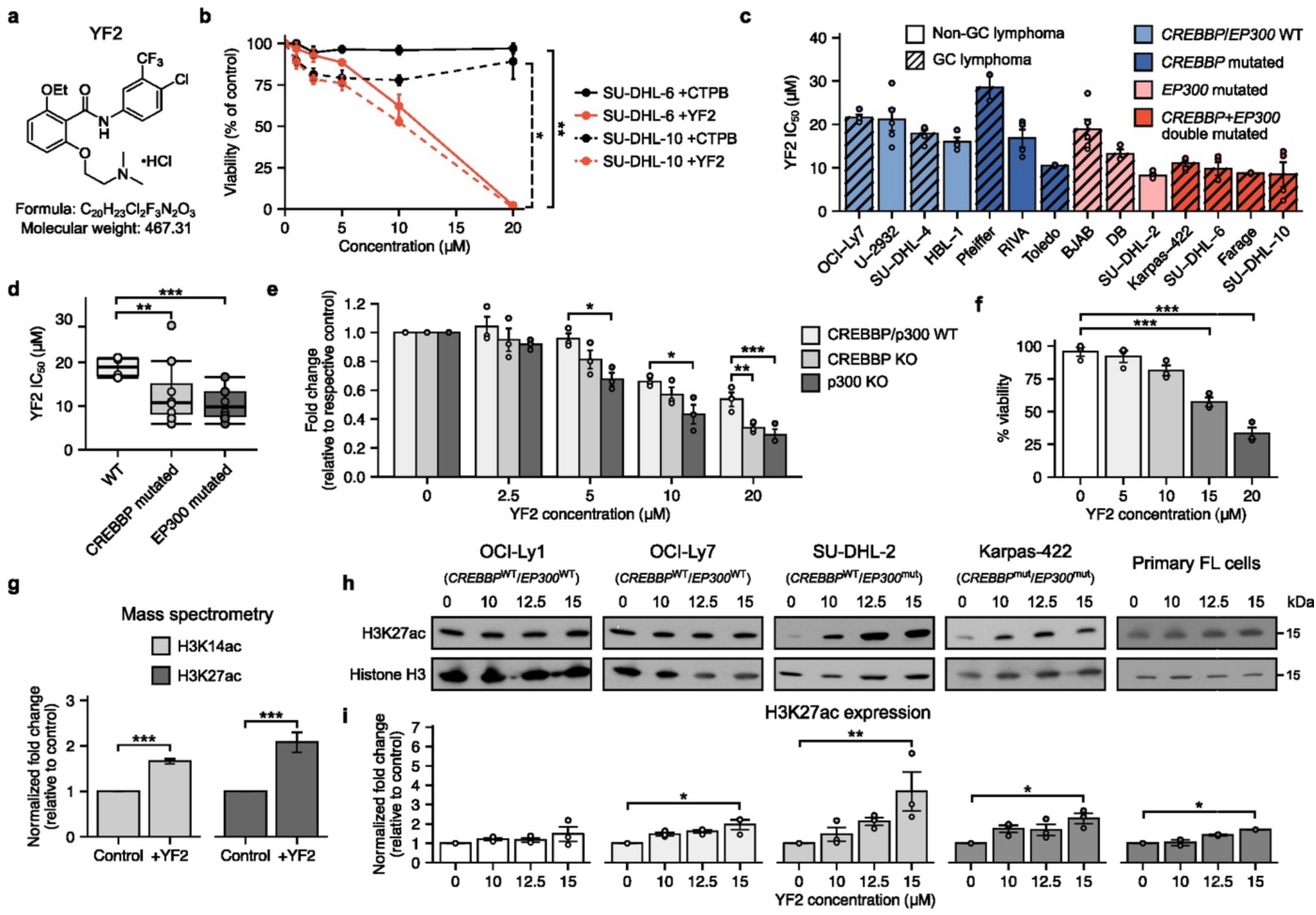
YF2 induces cytotoxicity in B-cell lymphoma, exhibiting heightened eaects in *CREBBP/EP300*-deficient cells. **a**, Chemical structure, formula, and molecular weight of YF2. **b**, Cell viability following 6-day treatment with HAT activators CTPB (black) or YF2 (red) in B-cell lymphoma cell lines. Viability is expressed as a percentage of respective vehicle-treated controls and shown as mean ± standard error of the mean (s.e.m.). Comparisons within each cell line were performed using linear regression (*n* = 3). **c**, YF2 half-maximal inhibitory concentration (IC_50_) values across B-cell lymphoma cell lines stratified by *CREBBP*/*EP300* mutational status. Data are shown as mean ± s.e.m. (*n* = 2–5). **d**, Summary of YF2 IC_50_ values across cell line groups shown in **c**. Data are presented as means; box plots indicate interquartile range with median. Group means were compared using one-way ANOVA with Tukey’s HSD post hoc test (n = 4–7). **e**, Viability of OCI-Ly7 wild-type (WT) and CREBBP/p300 knockout (KO) cells following 6-day YF2 treatment. Data are normalized to vehicle-treated controls and shown as mean ± s.e.m. Statistical comparisons were performed using one-way ANOVA with Tukey’s HSD post hoc test (*n* = 3). **f**, Viability of primary follicular lymphoma (FL) cells treated with YF2 for 6 days in vitro. Viable cells are expressed as a percentage of total cells and shown as mean ± s.e.m. Statistical comparisons were performed using one-way ANOVA with Tukey’s HSD post hoc test (*n* = 3). **g**, H3K14 and H3K27 acetylation in SU-DHL-6 cells treated with 6 µM YF2 for 48 h, assessed by mass spectrometry. Data are presented as fold change relative to vehicle-treated controls (mean ± s.e.m.) and analyzed using unpaired two-sided t-tests (*n* = 3). **h**, Immunoblot analysis of histone acetylation in B-cell lymphoma cell lines and primary FL cells treated with YF2 for 3 days (*n* = 3). **i**, Quantification of H3K27ac from **h**. Data are normalized to vehicle controls and shown as mean ± s.e.m. Statistical comparisons were performed using the Kruskal–Wallis test with Dunn’s post hoc correction (*n* = 3). Asterisks (*) denote statistical significance: * = *P* < 0.05, ** = *P* < 0.01, and *** = *P* < 0.001.

To evaluate YF2 sensitivity more broadly, the YF2 half-maximal inhibitory concentration (IC_50_) was determined across a diverse panel of B-cell lymphoma cell lines with various *CREBBP*/*EP300* mutations. *CREBBP*/*EP300* wild type (WT) cell lines were less sensitive to YF2 than those harboring *CREBBP* or *EP300* mutations (Figs. 1c,d). The copy number status and transcriptional expression of *CREBBP*+*EP300* in these cell lines, obtained from DepMap^27^, were largely consistent with sensitivity to YF2 (Supplementary Figs. S2a–c). In contrast, cell doubling time showed no correlation with YF2 IC_50_ values (Supplementary Figs. S2d–f), and no significant differences in YF2 sensitivity were observed between GC and non-GC B-cell lines (Supplementary Fig. S2g).

To directly test whether disruption of HAT signaling alters YF2 sensitivity, CREBBP and p300 were knocked out (KO; Supplementary Fig. S3a) in OCI-Ly7 cells, which are normally CREBBP/p300 WT, and then treated with increasing doses of YF2. Both KO cell lines exhibited increased sensitivity to YF2 compared to parental controls, confirming that altered CREBBP/p300 signaling enhances susceptibility (Fig. 1e). To extend these findings to primary samples, ascites fluid from a patient with FL was collected, and kappa light chain–restricted putative lymphoma cells were isolated by flow cytometric sorting (Supplementary Fig. S3b) before *in vitro* treatment with YF2. YF2 exposure reduced the viability of primary FL cells at higher concentrations (Fig. 1f).

Given the observed selective cytotoxicity, we next asked whether YF2 modulates histone acetylation in lymphoma cells. SU-DHL-6 cells were treated with YF2 *in vitro* and analyzed by mass spectrometry. YF2 exposure increased H3K14 and H3K27 acetylation compared to vehicle-treated controls (Fig. 1g). Consistent with these findings, total H3K27 acetylation was generally increased across GCB lymphoma cell lines and primary FL cells following YF2 treatment, as measured by immunoblotting (Figs. 1h,i).

### YF2 engages the CREBBP/p300 bromodomain to stimulate HAT activity and drive acetylation-dependent cell death

LBias, a computational tool for ligand-site prediction, was used to model YF2’s interaction with p300, predicting preferential binding to the bromodomain with possible interaction with the RING domain (Fig. 2a). Because the CREBBP/p300 RING domain can occlude the HAT substrate-binding pocket and restrict the auto-acetylation required for enzymatic activation^47,48^, YF2 binding may induce a conformational shift that relieves this autoinhibition, thereby promoting CREBBP/p300 auto-acetylation and HAT activity. To validate these predictions, surface plasmon resonance (SPR) was used to analyze YF2 binding to the p300 bromodomain in real time. Inobrodib, a known CREBBP/p300 bromodomain inhibitor, served as a positive control (Supplementary Figs. S4a,b). SPR analysis showed dose-dependent binding of YF2, with moderate kinetics reflected by the association rate constant (k_a_), rapid dissociation (k_d_), and low micromolar affinity (K_D_; Fig. 2b; Supplementary Figs. S4c,d). To further validate binding, thermal shift assays were performed to assess YF2’s effects on p300 bromodomain (Fig. 2c) and oT-target HAT domain binding (Fig. 2d). High YF2 concentrations increased melting temperature of the bromodomain region, consistent with an allosteric stabilization effect aligning with computational and SPR data.

**Fig. 2:**
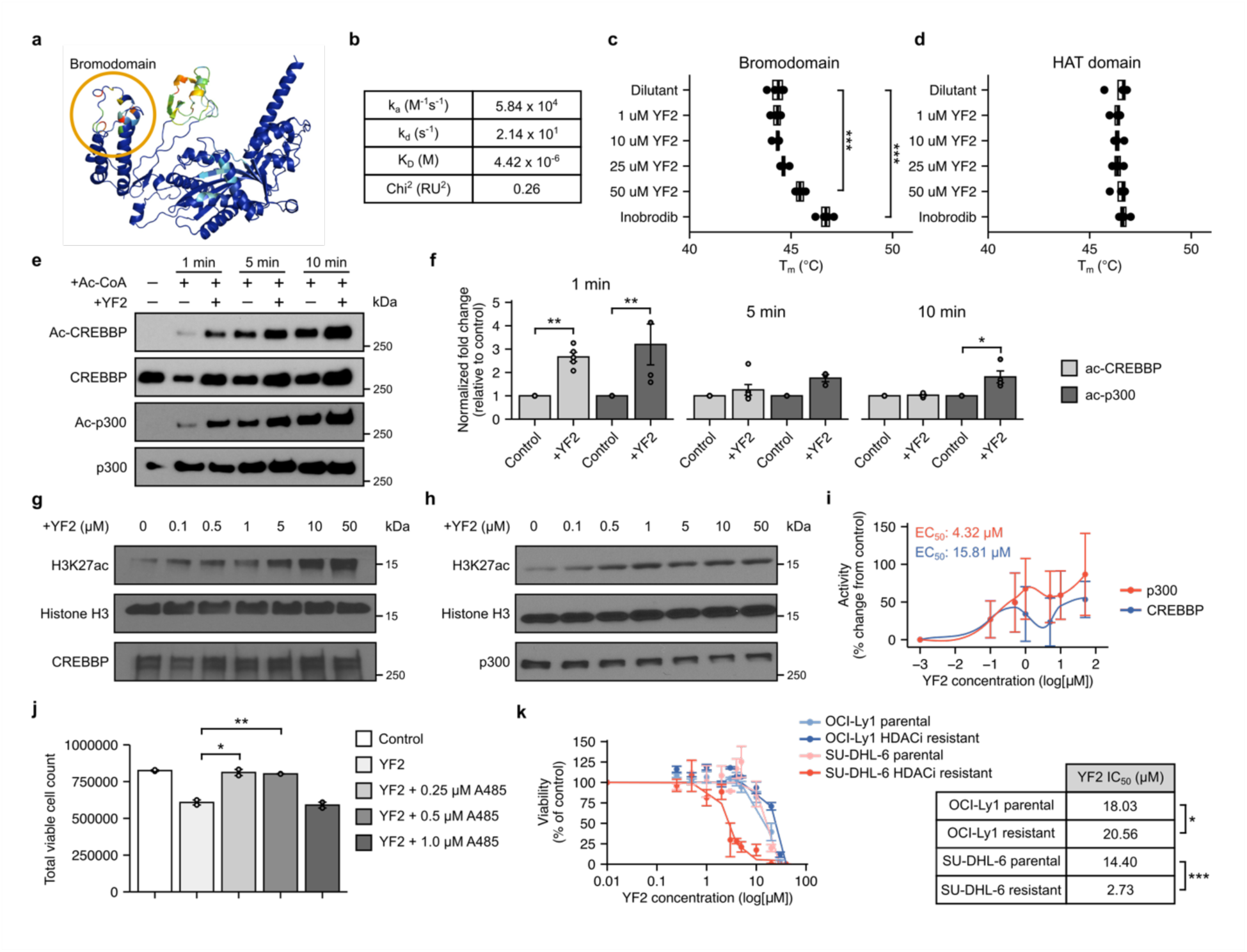
YF2 induces cytotoxicity in B-cell lymphoma, exhibiting heightened eaects in *CREBBP/EP300*-deficient conditions. **a**, Structural model of p300 showing predicted YF2 binding sites. The protein is shown in dark blue, with predicted interaction regions highlighted in warmer colors. **b**, Binding of YF2 to the p300 bromodomain assessed by surface plasmon resonance (SPR). Binding kinetics are described by the association rate constant (kₐ), dissociation rate constant (k_d_), and equilibrium dissociation constant (K_D_). **c**, Thermal shift analysis of YF2 binding to the p300 bromodomain. Inobrodib was used as a positive control. Data are presented as mean melting temperature (T_m_). Box plots indicate the interquartile range with the median. Statistical comparisons were performed using one-way ANOVA with Tukey’s HSD post hoc test (*n* = 4). **d**, Thermal shift analysis of YF2 binding to the p300 HAT domain. Data are presented as mean T_m_ with box plots indicating the interquartile range and the median. Statistical comparisons were performed using one-way ANOVA with Tukey’s HSD post hoc test (*n* = 4). **e**, Immunoblot analysis of CREBBP and p300 auto-acetylation following incubation with 50 nM YF2 for 1, 5, or 10 min (*n* = 3–6). **f**, Quantification of auto-acetylation shown in **e**. Data are normalized to vehicle-treated controls and presented as mean ± s.e.m. Comparisons at each time point were performed using the Mann–Whitney U test (*n* = 3–6). **g**, Cell-free H3K27 acetylation by CREBBP following 30 min exposure to increasing concentrations of YF2 (*n* = 3). **h**, Cell-free H3K27 acetylation by p300 following 30 min exposure to increasing concentrations of YF2 (*n* = 3). **i**, Quantification of H3K27 acetylation from **g** and **h**. Data are expressed as mean percentage change relative to vehicle-treated controls ± s.e.m., and EC_50_ values were determined by nonlinear regression (*n* = 3). **j**, Viability of OCI-Ly7 cells following 24 h treatment with YF2 in the presence or absence of the CREBBP/p300 inhibitor A-485. Data are presented as total viable cell counts (mean ± s.e.m.) and analyzed using one-way ANOVA with Tukey’s HSD post hoc test (*n* = 3). **k**, Viability of parental and HDAC inhibitor (HDACi)-resistant cell lines following 6-day treatment with increasing concentrations of YF2. Viability is expressed relative to vehicle-treated controls (mean ± s.e.m.), and IC_50_ values were compared using the Mann–Whitney U test (*n* = 3). Asterisks (*) denote statistical significance: * = *P* < 0.05 and ** = *P* < 0.01.

CREBBP and p300 bromodomains can recognize acetyl-lysine residues within their own structures, thereby regulating HAT activity^49^. To test whether YF2 modulates this autoregulatory mechanism, CREBBP and p300 auto-acetylation was assessed in a cell-free system at 1, 5, and 10 min. YF2 increased CREBBP auto-acetylation at 1 min and p300 auto-acetylation at 1 and 10 min (Figs. 2e,f). Increasing concentrations of YF2 also stimulated CREBBP- and p300-mediated H3K27 acetylation (Figs. 2g–i). In addition, YF2 enhanced H3 and H4 acetylation in fluorescent HAT assays (Supplementary Fig. S4f). Modest effects were also observed on KAT2A-dependent H4 acetylation (Supplementary Fig. S4g), while kinase profiling showed minimal oT-target activity (Supplementary Fig. S4e), suggesting relative selectivity for HAT enzymes. To confirm functional dependence on CREBBP/p300 activity, OCI-Ly7 cells were treated with YF2 and A-485, a CREBBP/p300 inhibitor. A-485 partially attenuated YF2-induced cytotoxicity, indicating dependency on HAT activation (Fig. 2j).

Finally, HDACi-resistant GCB lymphoma cell lines were generated (Supplementary Fig. S3c) and treated with YF2. While *CREBBP*/*EP300* WT HDACi-resistant OCI-Ly7 cells remained resistant to YF2, *CREBBP*+*EP300*-mutant SU-DHL-6 cells exhibited increased sensitivity compared to parental controls, suggesting that HAT activation may overcome epigenetic drug resistance (Fig. 2k).

### YF2 is well tolerated in vivo and improves overall survival in lymphoma mouse models

To evaluate pharmacokinetics *in vivo*, SCID/beige mice bearing SU-DHL-6 xenografts were treated with YF2 for five days (Fig. 3a). Both 40 and 60 mg/kg dosing achieved biologically relevant serum concentrations of YF2 (Fig. 3b), as well as measurable intratumoral drug levels (Figs. 3c,d). Tumor burden was significantly reduced in YF2-treated mice by day six compared to controls (Fig. 3e). Consistent with target engagement, tumors from treated mice showed increased H3K27 acetylation by mass spectrometry (Fig. 3f).

**Fig. 3:**
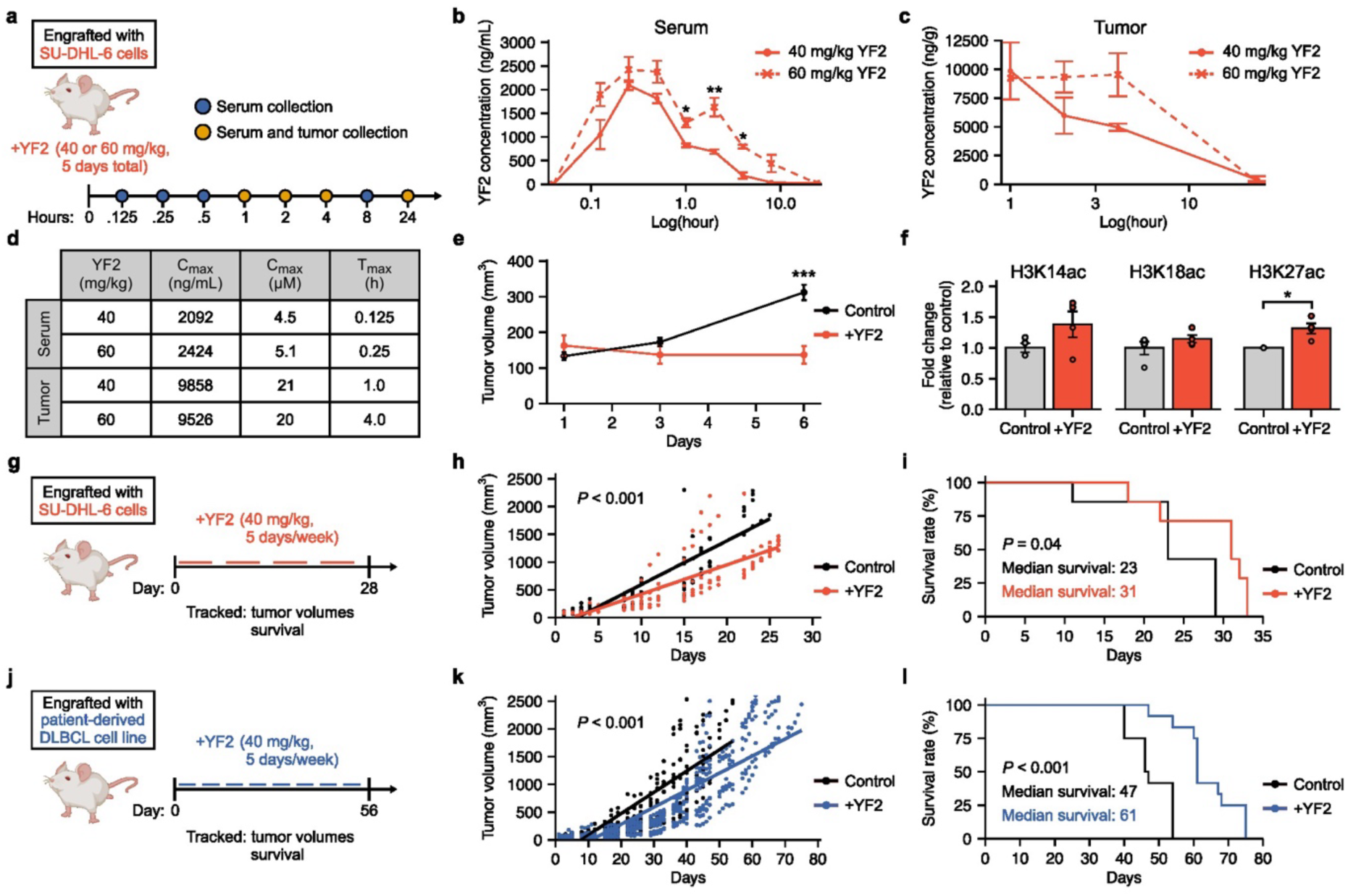
YF2 is well tolerated *in vivo* and improves overall survival in lymphoma mouse models. **a**, Experimental schematic for pharmacokinetic analysis. SCID/beige mice engrafted with SU-DHL-6 cells were treated with 40 or 60 mg/kg YF2 for 5 days, followed by serial serum and tumor collection over 24 h. Created with BioRender.com. **b**, YF2 concentrations in serum over 24 h following treatment. Data are shown as mean ± s.e.m., with comparisons at each time point performed using unpaired two-sided t-tests (*n* = 2–3). **c**, YF2 concentrations in tumors over 24 h following treatment. Data are shown as mean ± s.e.m., with comparisons at each time point performed using unpaired two-sided t-tests (*n* = 2–3). **d**, Maximum plasma concentration (C_max_) and time to maximum concentration (T_max_) derived from data in **b** and **c**. **e**, Tumor volumes in engrafted mice following treatment. Data are presented as mean ± s.e.m. and analyzed using linear regression (*n* = 4–8). **f**, Histone acetylation in tumors following 5-day treatment with 40 mg/kg YF2, assessed by mass spectrometry. Data are shown as fold change relative to vehicle-treated controls (mean ± s.e.m.) and analyzed using unpaired two-sided t-tests (*n* = 3). **g**, Experimental schematic for SU-DHL-6 xenograft study. Mice were treated with 40 mg/kg YF2 or vehicle for 28 days. Created with BioRender.com. **h**, Tumor volumes in SU-DHL-6 xenografts. Data are shown as individual values and analyzed using linear regression (*n* = 6–7). **i**, Kaplan–Meier survival analysis of SU-DHL-6 xenograft mice. Statistical comparisons were performed using a log-rank test (*n* = 7). **j**, Experimental schematic for patient-derived xenograft (PDX) model treated with YF2 or vehicle for 56 days. Created with BioRender.com. **k**, Tumor volumes in PDX xenografts. Data are shown as individual values and analyzed using linear regression (*n* = 12). **l**, Kaplan–Meier survival analysis of PDX xenograft mice. Statistical comparisons were performed using a log-rank test (*n* = 12). Asterisks (*) denote statistical significance: * = *P* < 0.05, ** = *P* < 0.01, and *** = *P* < 0.001.

To assess long-term efficacy, SU-DHL-6 xenograft-bearing mice were treated for 28 days (Fig. 3g). YF2 delayed tumor progression and improved survival compared to vehicle controls (Fig. 3h,i). These findings were extended to a patient-derived DLBCL xenograft model (Fig. 3j), where YF2 treatment for 56 days reduced tumor burden and prolonged survival relative to controls (Fig. 3k,l).

### YF2 enhances HAT-driven acetylation of p53, leading to changes in cell cycle progression and apoptosis

CREBBP and p300 acetylate both histone and non-histone substrates, including p53^11,12^. We therefore examined whether YF2 enhances p53 acetylation. Cell-free assays showed increased CREBBP-mediated (Fig. 4a) and p300-mediated (Fig. 4b) p53 acetylation following YF2 treatment (Fig. 4c). To account for frequent p53 mutations in lymphoma, functional p53 signaling was validated using an MDM2 inhibitor (Supplementary Fig. S5a). YF2 altered p53 acetylation and modulated downstream targets, including p21 and PUMA (Fig. 4d; Supplementary Figs. S5b–d).

**Fig. 4:**
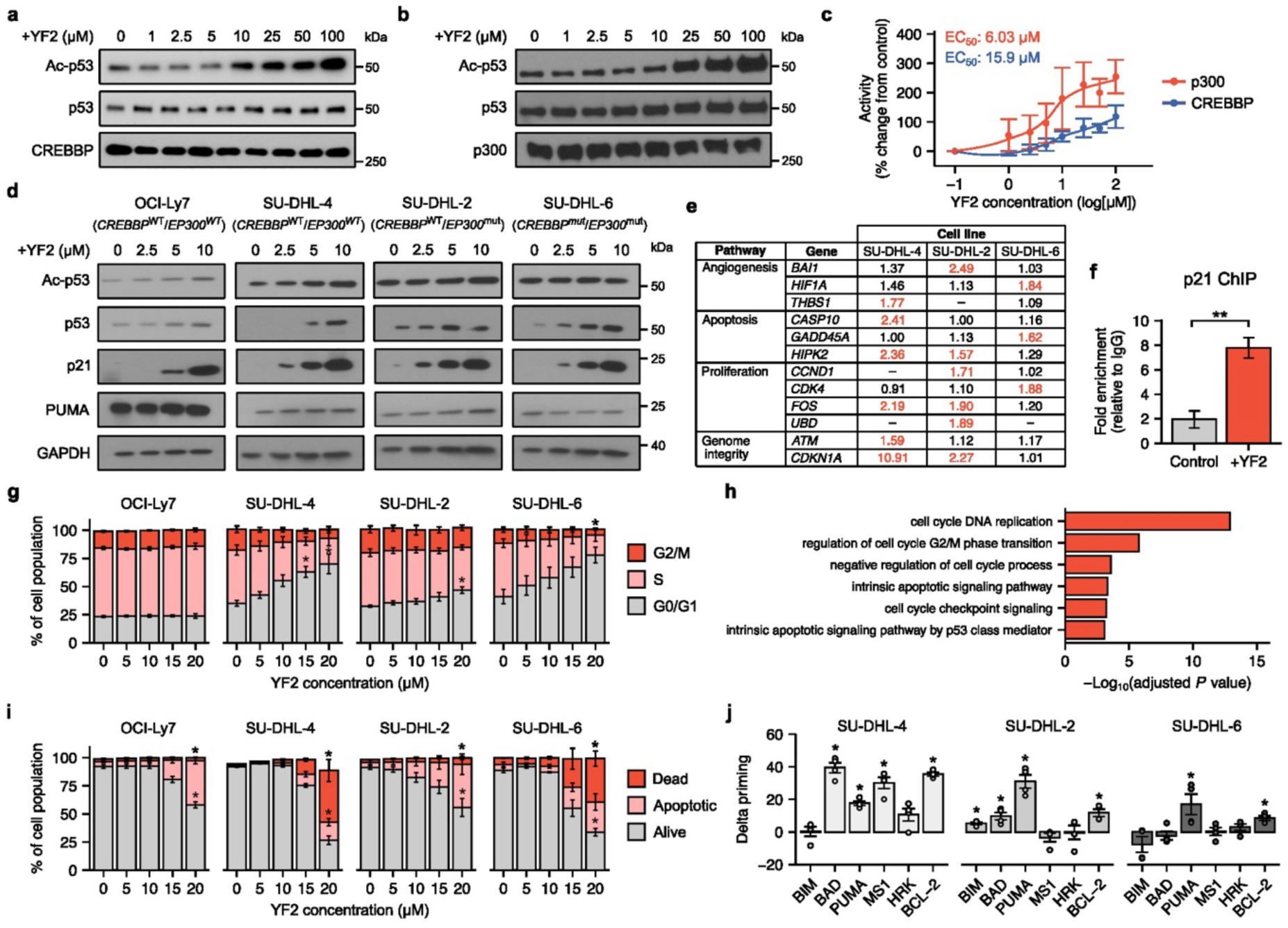
YF2 enhances HAT-driven acetylation of p53, leading to changes in cell cycle progression and apoptosis. **a**, Cell-free acetylation of p53 by CREBBP following 30 min exposure to increasing concentrations of YF2 (*n* = 3). **b**, Cell-free acetylation of p53 by p300 following 30 min exposure to increasing concentrations of YF2 (*n* = 3). **c**, Quantification of p53 acetylation from **a** and **b**. Data are expressed as percentage change relative to controls (mean ± s.e.m.), and EC_50_ values were calculated by nonlinear regression (*n* = 3). **d**, Immunoblot analysis of p53 and downstream targets in lymphoma cell lines treated with YF2 for 24 h (*n* = 3). **e**, Gene signaling array analysis of p53-associated genes following 24 h treatment with 10 µM YF2. Data are presented as fold change relative to controls; values > 1.5 are highlighted in red (*n* = 3). **f**, H3K27ac chromatin immunoprecipitation showing enrichment at the p21 locus following YF2 treatment. Data are presented as fold enrichment over IgG control (mean ± s.e.m.) and analyzed using unpaired two-sided t-tests (*n* = 3). **g**, Cell cycle distribution following 6-day YF2 treatment, assessed by flow cytometry. Data are shown as mean ± s.e.m. and analyzed using the Kruskal–Wallis test with Dunn’s post hoc correction (*n* = 3). **h**, Gene ontology over-representation analysis of up-regulated genes in YF2-treated SU-DHL-6 cells. Selected pathways with adjusted *P* < 0.05 are shown (*n* = 3–5). **i**, Apoptosis analysis following 6-day YF2 treatment, assessed by flow cytometry. Data are shown as mean ± s.e.m. and analyzed using the Kruskal–Wallis test with Dunn’s post hoc correction (*n* = 3). **j**, BH3 profiling following 3-day YF2 treatment. Data are shown as mean ± s.e.m. and analyzed using the Kruskal–Wallis test with Dunn’s post hoc correction (*n* = 3). Asterisks (*) denote statistical significance: * = *P* < 0.05 and ** = *P* < 0.01.

Gene signaling arrays further demonstrated upregulation of p53-associated pathways, including apoptosis, proliferation, and genome integrity (Fig. 4e). Chromatin immunoprecipitation confirmed increased H3K27ac at the p21 locus, consistent with transcriptional activation (Fig. 4f). Flow cytometry analysis showed that YF2 treatment increased the proportion of cells in G0/G1 and reduced G2/M, indicating cell cycle arrest (Fig. 4g; Supplementary Figs. S5e,f).

RNA-seq analysis in SU-DHL-6 (Supplementary Figs. S6a–d,f) and Karpas-422 cells (Supplementary Figs. 6g–k) revealed distinct transcriptional changes upon YF2 treatment. Pathway analysis showed enrichment of p53 signaling, apoptosis, and cell-cycle regulation (Fig. 4h; Supplementary Figs. S5f,l). Finally, functional assays demonstrated that YF2 increased apoptosis (Fig. 4i; Supplementary Fig. S5g,h) and enhanced mitochondrial priming for apoptosis via BH3 profiling (Fig. 4j).

### YF2 increases BCL6 acetylation, altering B-cell di:erentiation pathways and GC cycling

BCL6, a key regulator of GC biology and lymphoma pathogenesis^13^, is a known CREBBP/p300 substrate. YF2 increased CREBBP- (Fig. 5a) and p300-mediated (Fig. 5b) BCL6 acetylation in cell-free assays (Fig. 5c). To evaluate effects on GC biology *in vivo*, immunocompetent BALB/c mice were immunized and treated with YF2, followed by flow cytometric and spatial transcriptomic analysis of blood and spleen (Fig. 5d; Supplementary Figs. S7a,b). YF2 modestly reduced splenic mass relative to body weight (Supplementary Fig. S7c) and broadly altered B-cell maturation, decreasing immature B cells while increasing transitional and mature compartments (Fig. 5e). Within the GC compartment, YF2 increased dark zone GC B cells and reduced light zone populations, while also decreasing total splenic B cells and expanding plasma cell fractions (Fig. 5f–h). These shifts suggest altered GC cycling and enhanced differentiation toward terminal B-cell states.

**Fig. 5:**
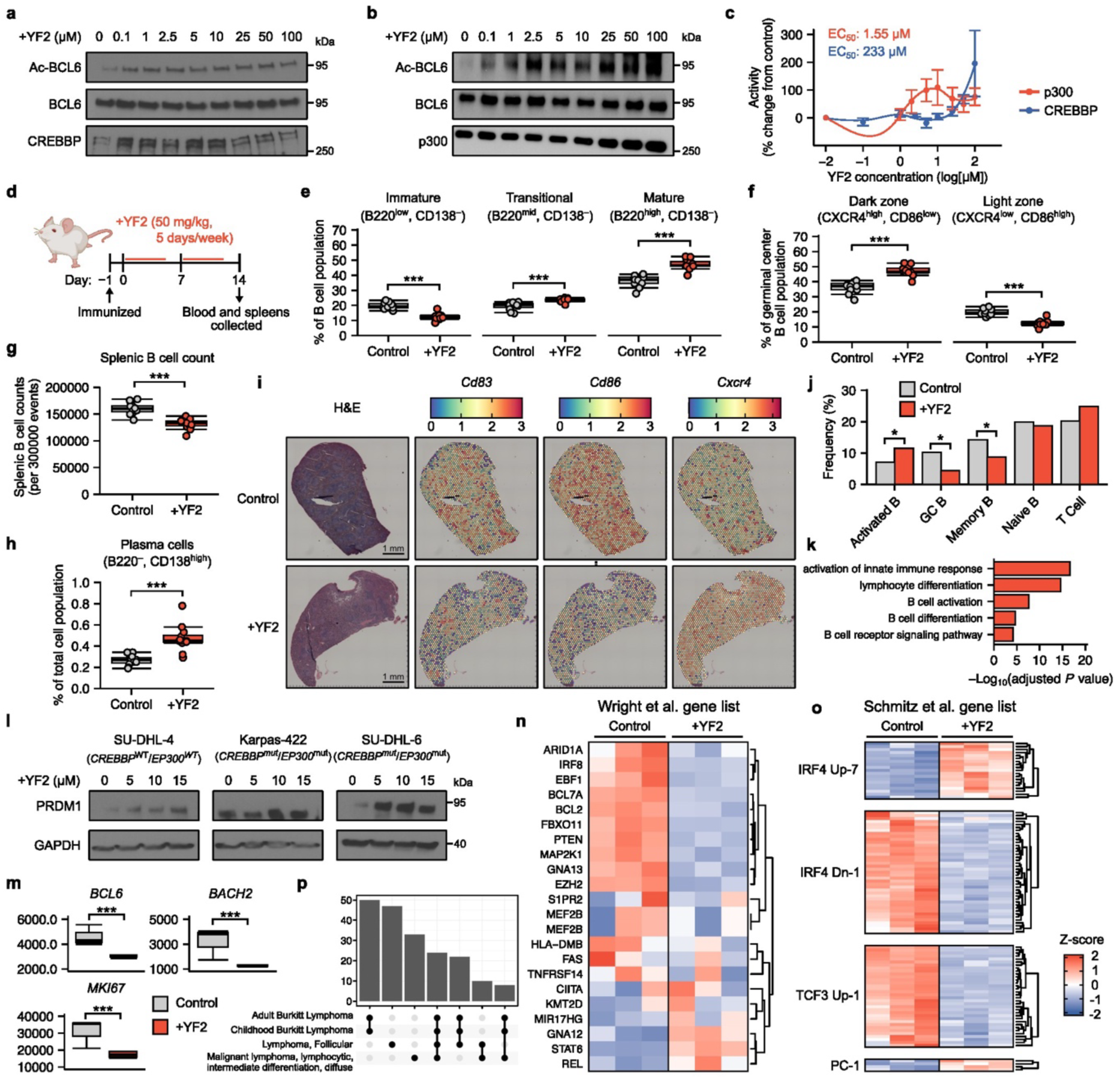
YF2 increases BCL6 acetylation, altering B-cell diaerentiation pathways. **a**, Cell-free acetylation of BCL6 by CREBBP following 30 min exposure to increasing concentrations of YF2 (*n* = 3). **b**, Cell-free acetylation of BCL6 by p300 following 30 min exposure to increasing concentrations of YF2 (*n* = 3). **c**, Quantification of BCL6 acetylation from **a** and **b**, expressed as percentage change relative to controls (mean ± s.e.m.); EC_50_ values were determined by nonlinear regression (*n* = 3). **d**, Experimental schematic of BALB/c mice treated with 50 mg/kg YF2 for 14 days before blood and spleen collection. Created with BioRender.com. **e**, B-cell differentiation states in spleen following treatment, expressed as a percentage of total B cells. Box plots indicate the interquartile range with the median. Statistical comparisons were performed using unpaired two-sided t-tests (*n* = 11–12). **f**, Germinal center dark and light zone populations following treatment, expressed as a percentage of GC B cells. Statistical comparisons were performed using unpaired two-sided t-tests (*n* = 11–12). **g**, Total splenic B-cell counts per 300,000 events. Statistical comparisons were performed using unpaired two-sided t-tests (*n* = 11–12). **h**, Plasma cell populations following treatment, expressed as a percentage of total cells. Statistical comparisons were performed using unpaired two-sided t-tests (*n* = 11–12). **i**, Spatial transcriptomic gene expression profiling in spleen following treatment (*n* = 1). **j**, Spatial transcriptomic clustering and cell-type composition by treatment condition. Differential abundance was assessed using Fisher’s exact test with Benjamini–Hochberg correction (*n* = 1). **k**, Gene ontology analysis of differentially expressed genes in the activated B-cell cluster identified in **j** (*n* = 1). **l**, PRDM1 protein expression following YF2 treatment, assessed by immunoblotting (*n* = 3). **m**, RNA-seq analysis of B-cell differentiation genes in SU-DHL-6 cells. Box plots indicate the interquartile range with the median (*n* = 3–5). **n**, Heatmaps showing YF2-induced changes in EZB-associated gene signatures defined by Wright et al.^31^ (*n* = 3–5). **o**, Heatmaps showing YF2-induced changes in EZB-associated gene signatures defined by Schmitz et al.^32^ (*n* = 3–5). **p**, Gene–disease association enrichment analysis of differentially expressed genes following YF2 treatment (*n* = 3–5). Asterisks (*) denote statistical significance: * = *P* < 0.05 and *** = *P* < 0.001.

Spatial transcriptomic analysis of spleens (Supplementary Figs. S8a,b) showed decreased expression of light zone markers *Cd83* and *Cd86* and increased expression of the dark zone marker *Cxcr4* (Fig. 5i; Supplementary Fig. S8c). Unsupervised clustering and cell annotation identified multiple immune populations, including GC B-cell subsets, with YF2 altering the abundance of activated, GC, and memory B-cell states (Fig. 5j; Supplementary Figs. S8d–k). Gene ontology analysis of activated B cells indicated enrichment of differentiation and activation programs (Fig. 5k). The spatial transcriptomic findings indicating altered GC cycling and B-cell differentiation were further supported by immunohistochemistry (Supplementary Figs. S9a–c).

In lymphoma cell lines, YF2 increased PRDM1 expression (Fig. 5l; Supplementary Fig. S7d) and reduced expression of BCL6 and BACH2, consistent with promotion of terminal differentiation (Fig. 5m; Supplementary Fig. S7e). RNA-seq further showed reversal of EZB subtype-associated transcriptional signatures described by Wright et al. ^50^ and Schmitz et al.^51^ (Figs. 5n,o; Supplementary Figs. S6l,m). Gene-disease enrichment analysis confirmed a strong association of YF2-responsive genes with lymphoma-related pathways (Fig. 5p).

### YF2 increases markers of antigen presentation and improves immune checkpoint blockade therapy

Given its effects on B-cell differentiation, we next assessed whether YF2 alters antigen presentation pathways. Gene expression profiling of SU-DHL-6 cells showed up-regulation of multiple MHC-related genes (Fig. 6a). Flow cytometry confirmed increased MHC-I and MHC-II surface expression following YF2 treatment (Figs. 6b,c), and immunoblotting supported these findings (Supplementary Figs. S10a,b). Chromatin immunoprecipitation further showed enrichment of antigen-presentation genes (Supplementary Fig. S10c). Gene set enrichment analysis confirmed activation of antigen processing and presentation genes and pathways (Fig. 6d; Supplementary Figs. 6e). *In vivo*, immunized BALB/c mice treated with YF2 exhibited increased MHC-I and MHC-II expression on peripheral B cells (Figs. 6e,f). YF2 also increased circulating CD8^+^ T-cell proportions without affecting PD-1^+^ exhausted subsets (Fig. 6g; Supplementary Fig. S10g), and elevated PD-L1 expression on B cells (Supplementary Fig. S10d).

**Fig. 6:**
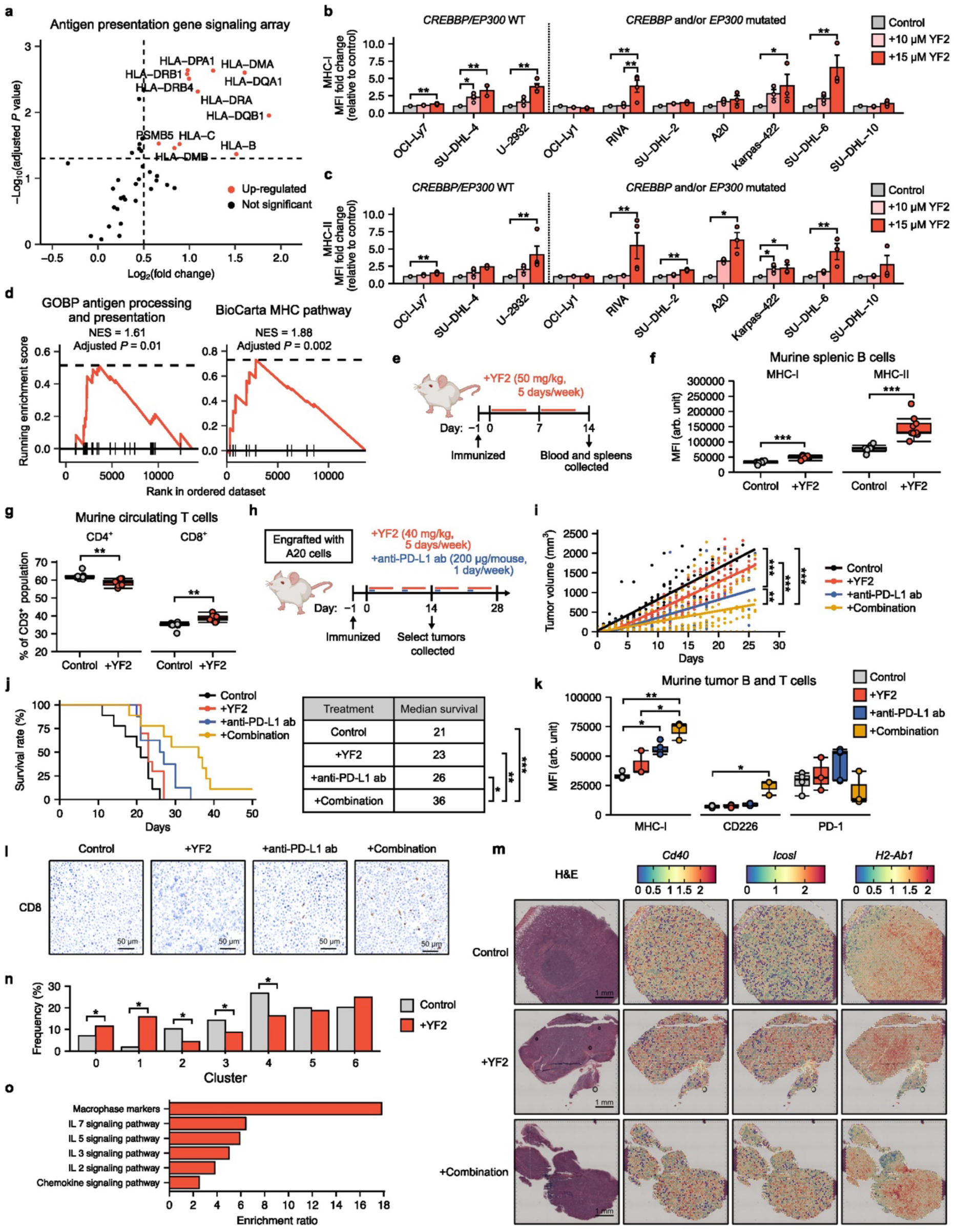
YF2 increases markers of antigen presentation and improves immune checkpoint blockade therapy. **a**, Gene signaling array showing differential expression of antigen presentation genes following 24 h treatment with 10 µM YF2. Genes with log₂(fold change) > 0.5 and adjusted *P* < 0.05 are highlighted (*n* = 3). **b**, Surface expression of MHC-I in lymphoma cell lines following 6-day YF2 treatment, assessed by flow cytometry. Data are shown as fold change relative to controls (mean ± s.e.m.) and analyzed using the Kruskal–Wallis test with Dunn’s post hoc correction (*n* = 3). **c**, Surface expression of MHC-II in lymphoma cell lines following 6-day YF2 treatment, assessed by flow cytometry. Data are shown as fold change relative to controls (mean ± s.e.m.) and analyzed using the Kruskal–Wallis test with Dunn’s post hoc correction (*n* = 3). **d**, Gene set enrichment analysis showing activation of antigen processing and presentation pathways following YF2 treatment (*n* = 3–5). **e**, Experimental schematic of BALB/c mice treated with YF2 for 14 days prior to blood collection. Created with BioRender.com. **f**, MHC-I and MHC-II expression on peripheral B cells following treatment, measured as mean fluorescence intensity (MFI). Box plots indicate the interquartile range with the median. Statistical comparisons were performed using unpaired two-sided t-tests (*n* = 9–10). **g**, CD4^+^ and CD8^+^ T-cell populations in peripheral blood following treatment. Box plots indicate the interquartile range with the median. Statistical comparisons were performed using unpaired two-sided t-tests (*n* = 9–10). **h**, Experimental schematic of combination therapy study with YF2 and anti–PD-L1 antibody in A20 lymphoma-bearing mice. Created with BioRender.com. **i**, Tumor volumes in A20 xenografts. Data are shown as individual values and analyzed using linear regression (*n* = 9–10). **j**, Kaplan–Meier survival analysis of A20 xenograft mice. Statistical comparisons were performed using a log-rank test (*n* = 9–10). **k**, Expression of immune activation markers in tumors, assessed by flow cytometry. Data are shown as MFI (mean ± s.e.m.) and analyzed using the Kruskal–Wallis test with Dunn’s post hoc correction (*n* = 3–5). **l**, Immunohistochemical analysis of intratumoral CD8^+^ T cells (*n* = 3). **m**, Spatial transcriptomic profiling of immune-related gene expression across treatment groups (*n* = 1). **n**, Spatial cluster composition of tumor samples by treatment (*n* = 1). **o**, Pathway enrichment analysis of up-regulated genes in cluster 1 identified in **n** (*n* = 1). Asterisks (*) denote statistical significance: * = *P* < 0.05, ** = *P* < 0.01, and *** = *P* < 0.001.

To test therapeutic synergy, BALB/c mice bearing A20 lymphoma were treated with YF2 and anti–PD-L1 antibody (Fig. 6h; Supplementary Figs. S10e,f). Combination therapy significantly reduced tumor burden and improved survival compared to either monotherapy (Fig. 6i,j) with no effect and mouse weights (Supplementary Fig. S10h). Tumor analysis revealed increased MHC-I and CD226 expression, consistent with enhanced cytotoxic immune activity (Fig. 6k), as well as increased CD8^+^ infiltration by immunohistochemistry (Fig. 6l; Supplementary Fig. S10i).

Spatial transcriptomic profiling of tumors showed increased expression of antigen presentation and immune activation genes, including *Cd40*, *Icosl*, and *H2-Ab1*, in YF2- and combination-treated tumors (Fig. 6m; Supplementary Figs. S11a,b). Unsupervised clustering revealed treatment-dependent shifts in tumor microenvironment composition, particularly in cluster 1 (Fig. 6n), which showed enrichment of cytokine signaling and T-cell activation pathways (Fig. 6o). Deconvolution analysis further identified a putative T-cell population (topic 3) with increased immune-response gene signatures in YF2-treated tumors (Supplementary Figs. S11c–h; Supplementary Data 3,4).

## Discussion

Among patients diagnosed with GCB lymphomas and treated with standard therapies, 30% experience relapse, and fewer than half are eligible for intensive salvage treatments such as CAR T-cell therapy^52^. While many newly developed therapies are largely agnostic to the biological drivers of GCB lymphoma or show greater efficacy in non-GCB subtypes, recent advances in molecular pathogenesis have renewed interest in precision-targeted approaches. In particular, epigenetic derangements arising from mutations in histone-modifying proteins have emerged as key drivers of lymphomagenesis^53^. Previous studies have shown that monoallelic mutations in HAT enzymes contribute to tumor suppressor silencing; however, whether pharmacological activation of the remaining functional allele or an unaffected paralogue can reverse the epigenetic dysregulation in GCB malignancies remains unclear. Here, we show that HAT activation induces cytotoxicity in malignant B cells and partially reverses lymphomagenesis-associated transcriptional programs, supporting its therapeutic potential in GCB lymphoma.

Beyond tumor-intrinsic effects, YF2 may also counteract immune evasion driven by epigenetic alterations^54,55^. Loss of function in CREBBP/p300 leads to repression of MHC-II expression, which is required for antigen presentation, B-cell maturation, and plasma cell differentiation^56^, and reduced MHC-II expression is associated with impaired tumor immunosurveillance and poor outcomes in DLBCL^57^. Consistent with this, YF2 increases MHC-I and MHC-II expression and activates antigen presentation pathways, suggesting restoration of tumor immune visibility. These findings support a role for epigenetic therapies in enhancing immunotherapy responses^58–60^. Anti-PD-L1 antibodies and other immune checkpoint blockade inhibitors show promise in multiple malignancies^61,62^ but have limited single-agent efficacy in GCB lymphoma^63^. In this context, combining YF2 with anti-PD-L1 reduces tumor burden and improves survival compared to either agent alone, while increasing intratumoral T-cell activity. Together, these results support HAT activation as a strategy to enhance responses to immune checkpoint blockade in GCB lymphoma.

A limitation of this study is the treatment of *CREBBP* and *EP300* mutations as functionally equivalent epigenetic derangements. CREBBP and p300 share many targets and exhibit partial redundancy^64^, and activity from either acetyltransferase is required for B-cell development and survival^65^. However, they also regulate distinct transcriptional programs in GC subcompartments^66^. Mutational inactivation of CREBBP, but not p300, promotes tumor growth in xenograft models^15^, whereas deletion of p300, but not CREBBP, reduces the expression of genes involved in cell cycle progression and DNA replication in GC B cells^66^. The frequency and type of CREBBP and p300 mutations also differ considerably among GCB lymphoma subtypes and may affect their susceptibility to HAT activation^67^. YF2 effectively activates both CREBBP and p300, but the relative contribution of each enzyme to the therapeutic response was not assessed.

Epigenetic therapies generally show modest single-agent activity in relapsed or refractory GCB lymphoma^68,69^, whereas combination approaches demonstrate more robust and synergistic effects in preclinical models^70^. Given the clustering of epigenetic mutations in GC-derived lymphomas^50,51^, dual targeting of complementary epigenetic pathways may therefore be advantageous. EZH2, a histone methyltransferase frequently mutated in GCB lymphoma, is associated with reduced MHC-II expression and immune evasion^71^. Mutations in *CREBBP*/*EP300* and *EZH2* may cooperate to suppress antigen presentation and T-cell activation, suggesting that combined HAT activation and EZH2 inhibition could be synergistic. Further studies are needed to evaluate YF2 in combination with other epigenetic therapies.

In summary, YF2 induces cytotoxicity in lymphoma cells, is well tolerated *in vivo*, and improves survival both as a monotherapy and in combination with immunotherapy in lymphoma mouse models. Collectively, these findings support HAT activation as a promising therapeutic strategy and highlight epigenetic reprogramming as a means to enhance immunotherapy efficacy in GCB lymphomas.

## Supporting information

File containing all supplemental figures

## Acknowledgments

Research reported in this publication was supported by the National Cancer Institute (NCI) of the National Institutes of Health (NIH) under grant R01CA222931 (J.E.A). J.E.A was also supported by Appia Pharmaceuticals, the Irving Scholars Program of the Columbia University Irving Institute for Clinical and Translational Research, and The Esther and Oded Aboodi Lymphoma Research Fund. Y.L. was sponsored by the Lymphoma Research Foundation Postdoctoral Fellowship Grant. B.H. was supported by NIH grant R35GM139585. Research was performed at Columbia University by the CCTI Flow Cytometry Core, supported by NIH grant S10OD030282, the Genomics and High Throughput Screening Shared Resource, supported by NIH grant P30CA013696, the Molecular Pathology Shared Resource, supported by NIH grant P30CA013696, and the Biomarkers Core Laboratory, supported by NIH grant UL1TR001873. Proteomics services were performed by the Northwestern Proteomics Core Facility, supported by NCI grant P30CA060553 and NIH grant S10OD025194, and the National Resource for Translational and Developmental Proteomics, supported by grant P41GM108569. The content of this paper is solely the responsibility of the authors and does not necessarily represent the official views of the NIH.

All compounds in the RA series are proprietary to Appia Pharmaceutical and are protected by WO2020163731. The remaining compounds are protected through WO2011072243, WO2012088420, WO2018017858, and WO2015153410, and have been licensed by Columbia University to Appia Pharmaceuticals.

## Author Contributions

T.B.P., Y.L., B.E., and J.E.A. conceived the project, T.B.P., Y.L., B.E., E.C.R., M.P., Y.G., S.S.T., and Y.K.R.T. conducted general experiments, C.K. performed the medium throughput screening, S.C. and R.N. completed the pharmacokinetic analyses, B.H. and H.H. performed the computational modeling, N.L.K., J.C., and N.A.A. conducted the mass spectrometry, T.B.P., A.O., and C.L analyzed the spatial transcriptomic data, T.B.P. ran the statistical analyses, J.E.A. supervised the research and provided funding, and T.B.P. and J.E.A. wrote the manuscript with input from all authors.

## Data Availability

Bulk RNA-seq and spatial transcriptomic files have been deposited in the National Center for Biotechnology Information Gene Expression Omnibus database under accession numbers GSE294618 and GSE295100, respectively. All other materials are available from the corresponding author upon request.

