## Supplementary material for "A Novel CREBBP/p300 Activator, YF2, Enhances Cytotoxic and Immune-Mediated Responses in B-Cell Lymphoma": File containing all supplemental figures

[1]Division of Hematology and Oncology, Department of Medicine, Herbert Irving Comprehensive Cancer Center, Columbia University, New York, NY, USA
[2]Division of Hematology, Oncology, Stem Cell Transplant, and Regenerative Medicine, Department of Pediatrics, Stanford University, Stanford, CA, USA
[3]Division of Hematology, Mayo Clinic, Rochester, MN, USA
[4]Department of Medicine, Columbia University Irving Medical Center, New York, NY, USA
[5]Department of Systems Biology, Columbia University, New York, NY, USA
[6]Department of Pathology and Cell Biology, Columbia University, New York, NY, USA
[7]Irving Institute for Clinical and Translational Research, Columbia University, New York, NY, USA
[8]Department of Biochemistry and Molecular Biophysics, Columbia University, New York, NY, USA
[9]Department of Chemistry, Molecular Biosciences, and the National Resource for Translational and Developmental Proteomics, Northwestern University, Evanston, IL, USA
[10]Departments of Chemistry and Molecular Biosciences and the Feinberg School of Medicine, Northwestern University, Chicago, IL, USA
*These authors contributed equally
[†]Corresponding author

**Description of Supplementary Files**

File Name: Supplementary Data 1
Description: Differential gene expression analysis comparing YF2- and vehicle-treated SU-DHL-6 RNA-seq samples.

File Name: Supplementary Data 2
Description: Whole-exome sequencing data and SNP/indel analysis of the A20 mouse cell line.

File Name: Supplementary Data 3
Description: Gene expression profiles for all deconvoluted spatial topics in the vehicle-treated Visium tumor sample.

File Name: Supplementary Data 4
Description: Gene expression profiles for all deconvoluted spatial topics in the YF2-treated Visium tumor sample.

**a**

| | SU-DHL-10 | | SU-DHL-2 | Farage | | HBL-1 | |
|---|---|---|---|---|---|---|---|
| Compound | 72 h | 144 h | 72 h | 72 h | 144 h | 72 h | 144 h |
| YF2 | 20.523 | 30.012 | 7.5216 | 13.716 | 9.5373 | 10.371 | 6.2275 |
| JF1 | 6.2502 | 6.955 | 3.6653 | 4.8147 | 4.6602 | 5.2524 | 3.6772 |
| JF3 | 17.324 | 19.632 | 13.23 | no curve | 11.289 | no curve | no curve |
| JF4 | no curve | 12.68 | 15.005 | no curve | no curve | no curve | no curve |
| JF7 | no curve | no curve | 11.354 | 148.7 | 22.79 | 17.671 | 11.359 |
| JF10 | 7.9634 | 9.9566 | 4.8084 | 4.8473 | 4.2839 | 4.0022 | 4.6553 |
| JF16 | no curve | no curve | no curve | 44.408 | no curve | no curve | no curve |
| RA010115 | 20.498 | no curve | no curve | no curve | no curve | no curve | no curve |
| RA010143 | 14.028 | 14.119 | 4.7153 | 7.0379 | 6.0122 | 7.5616 | 6.1592 |
| RA010146 | no curve | 865.34 | no curve | no curve | no curve | 11.364 | 8.4014 |
| RA010150 | no curve | no curve | no curve | 178.2 | 150 | no curve | 96.742 |
| RA010160 | 18.519 | 21.44 | no curve | no curve | 851.53 | 19.458 | 17.062 |
| RA010165 | 11.738 | 13.447 | 8.1497 | 10.746 | 10.505 | 10.544 | 9.0911 |
| RA010166 | 18.133 | 14.208 | no curve | no curve | no curve | no curve | no curve |
| RA010168 | no curve | no curve | no curve | 65.052 | 104.04 | no curve | no curve |
| RA010171 | no curve | no curve | no curve | no curve | no curve | 16467 | 60.504 |
| RA010900 | no curve | 148.12 | 36.57 | 19.062 | 28.406 | 38.78 | 16.266 |
| RA013005 | 24.395 | 23.122 | 18.661 | 23.93 | 13.903 | 13.853 | 11.023 |
| RA013886 | 21.568 | 93.257 | 54.72 | 19.142 | 21.699 | 21.278 | 15.813 |
| RA013894 | no curve | 14.437 | 7.5431 | 8.0899 | 6.9111 | 6.2805 | 4.3382 |
| RA013895 | 11.276 | 11.713 | 9.7473 | 11.277 | 10.32 | 10.142 | 6.8794 |
| RA013905 | 30.456 | 31.073 | 28.58 | 62.8 | no curve | no curve | 43.218 |
| RA013911 | 15.689 | 14.915 | 5.7254 | 8.8349 | 6.7775 | 9.5275 | 5.2325 |
| RA012919 | 20.228 | 22.579 | 20.369 | 140.02 | 1213.8 | 5539.1 | 42.329 |
| RA013938 | 10.101 | 10.018 | 3.9755 | 5.6595 | 5.5705 | 4.8808 | 4.6546 |
| Romidepsin | 0.0011778 | 0.0011317 | 0.00054529 | 0.00094973 | 0.00059234 | 0.00057955 | 0.00040468 |
| RP14 | 8.3933 | 0.060207 | no curve | 73.943 | no curve | no curve | 64.406 |
| RP23 | 10.594 | 7.6572 | 12.478 | 5.9447 | 7.6717 | 13.557 | 4.5456 |
| RP58 | no curve | no curve | no curve | 56.248 | no curve | no curve | 64.406 |
| RP59 | 9.7506 | 10.083 | 6.453 | 9.2502 | 8.27 | 3.415 | 4.9561 |
| Thimerosal | 0.05405 | 0.045239 | 0.36321 | 0.074302 | 0.082608 | 0.071289 | 0.074068 |

**b**

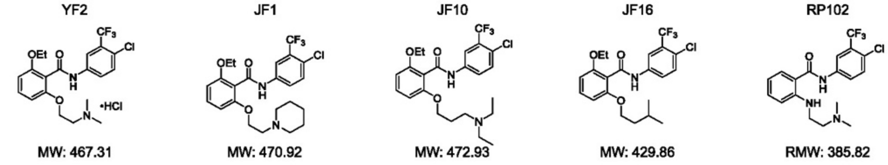


**c**

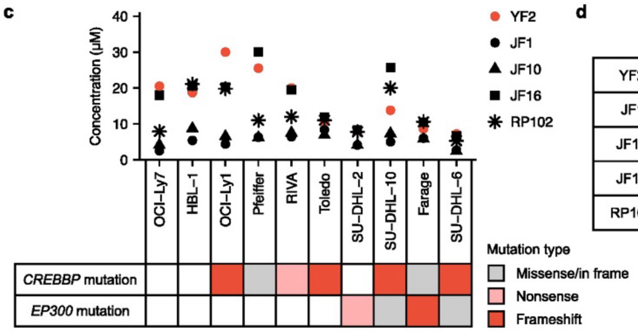


**d**

| | WT $IC_{50}$ | *CREBBP*$^{mut}$ $IC_{50}$ | *EP300*$^{mut}$ $IC_{50}$ | WT $IC_{50}$ / *CREBBP*$^{mut}$ $IC_{50}$ | WT $IC_{50}$ / *EP300*$^{mut}$ $IC_{50}$ |
|---|---|---|---|---|---|
| YF2 | 19.60 | 16.54 | 9.43 | 1.19 | 2.08 |
| JF1 | 3.95 | 5.60 | 4.45 | 0.71 | 0.89 |
| JF10 | 6.40 | 6.09 | 4.90 | 1.05 | 1.31 |
| JF16 | 19.25 | 17.79 | 12.78 | 1.08 | 1.51 |
| RP102 | 14.50 | 12.81 | 10.93 | 1.13 | 1.33 |

**Supplementary Fig. S1: Screening and selection of HAT-activating compounds in B-cell lymphoma.**

**a**, Medium-throughput cytotoxicity screening of HAT-activating compounds across four B-cell lymphoma cell lines. Data are presented as half-maximal effective concentrations ($EC_{50}$, μM; *n* = 3).

**b**, Chemical structures and molecular weights (MW) of five analog compounds sharing the N-phenyl benzamide scaffold selected for downstream analyses.

**c**, Half-maximal inhibitory concentrations ($IC_{50}$) of novel HAT-activating compounds across B-cell lymphoma cell lines harboring various *CREBBP*/*EP300* mutations. Data are presented as mean $IC_{50}$ values (*n* = 3).

**d**, Mean $IC_{50}$ values of each compound across wild-type (WT) and $CREBBP^{mut}/EP300^{mut}$ cell lines shown in **c**. Selective cytotoxicity is represented as the WT-to-mutant $IC_{50}$ ratio, with higher values indicating greater selectivity.

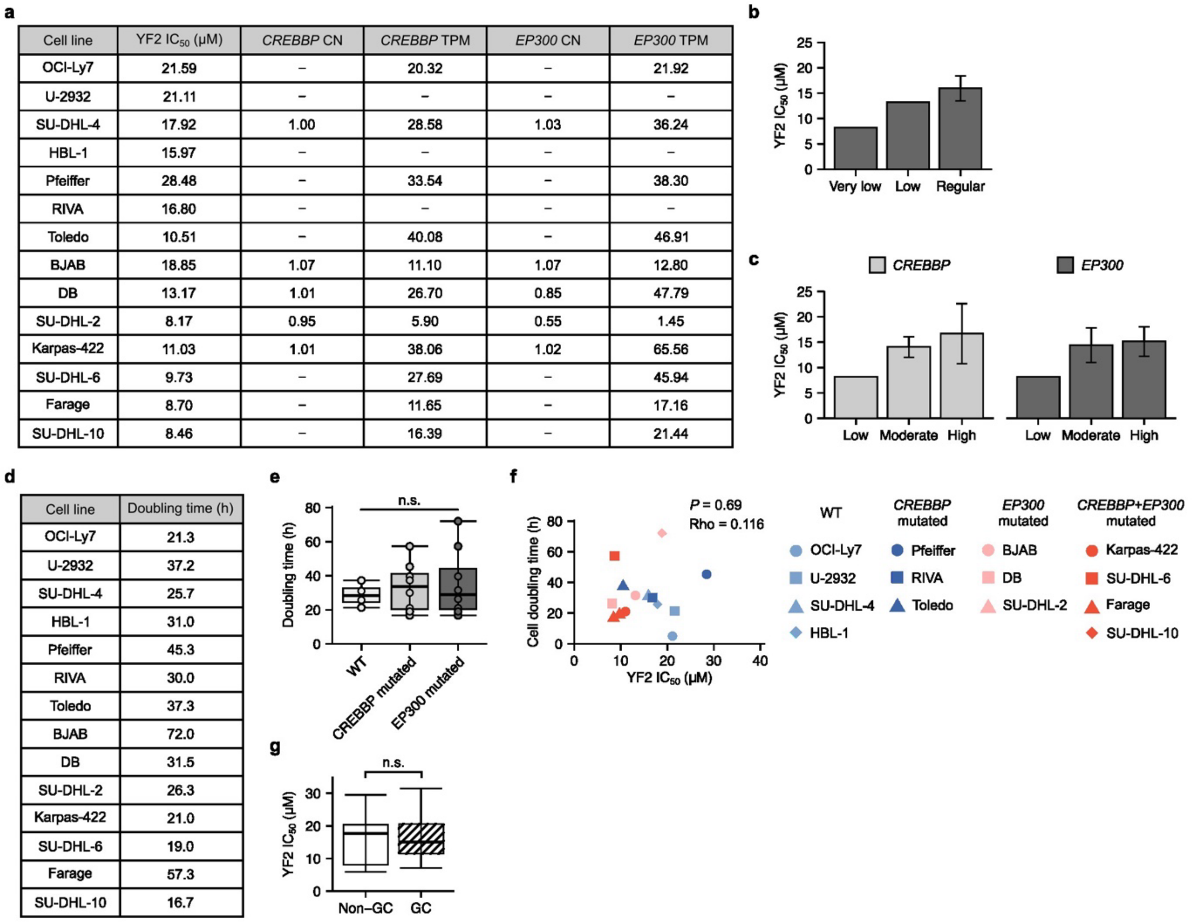


**a**

| Cell line | YF2 $IC_{50}$ (µM) | *CREBBP* CN | *CREBBP* TPM | *EP300* CN | *EP300* TPM |
|---|---|---|---|---|---|
| OCI-Ly7 | 21.59 | – | 20.32 | – | 21.92 |
| U-2932 | 21.11 | – | – | – | – |
| SU-DHL-4 | 17.92 | 1.00 | 28.58 | 1.03 | 36.24 |
| HBL-1 | 15.97 | – | – | – | – |
| Pfeiffer | 28.48 | – | 33.54 | – | 38.30 |
| RIVA | 16.80 | – | – | – | – |
| Toledo | 10.51 | – | 40.08 | – | 46.91 |
| BJAB | 18.85 | 1.07 | 11.10 | 1.07 | 12.80 |
| DB | 13.17 | 1.01 | 26.70 | 0.85 | 47.79 |
| SU-DHL-2 | 8.17 | 0.95 | 5.90 | 0.55 | 1.45 |
| Karpas-422 | 11.03 | 1.01 | 38.06 | 1.02 | 65.56 |
| SU-DHL-6 | 9.73 | – | 27.69 | – | 45.94 |
| Farage | 8.70 | – | 11.65 | – | 17.16 |
| SU-DHL-10 | 8.46 | – | 16.39 | – | 21.44 |

**d**

| Cell line | Doubling time (h) |
|---|---|
| OCI-Ly7 | 21.3 |
| U-2932 | 37.2 |
| SU-DHL-4 | 25.7 |
| HBL-1 | 31.0 |
| Pfeiffer | 45.3 |
| RIVA | 30.0 |
| Toledo | 37.3 |
| BJAB | 72.0 |
| DB | 31.5 |
| SU-DHL-2 | 26.3 |
| Karpas-422 | 21.0 |
| SU-DHL-6 | 19.0 |
| Farage | 57.3 |
| SU-DHL-10 | 16.7 |

**Supplementary Fig. S2: Relationship between YF2 sensitivity and genomic/cellular features.**

**a**, *CREBBP* and *EP300* copy number (CN) and transcript abundance expressed as transcripts per million (TPM) across lymphoma cell lines, obtained from the DepMap Portal.

**b**, Mean YF2 $IC_{50}$ across cell lines grouped by *CREBBP*/*EP300* CN status: regular (≥1.0), low (0.7–1.0), and very low (<0.7) ($n$ = 1–3).

**c**, Mean YF2 $IC_{50}$ across cell lines grouped by *CREBBP*/*EP300* TPM expression: high (>30), moderate (10–30), and low (<10) ($n$ = 1–7).

**d**, Cell doubling times of B-cell lymphoma cell lines ($n$ = 3).

**e**, Combined doubling times across cell types shown in **d**. Data are presented as means, with box plots indicating quartiles. Group comparisons were performed using one-way ANOVA with Tukey's HSD post-hoc test ($n$ = 4–8).

**f**, Correlation between cell doubling time and YF2 $IC_{50}$ values analyzed using Spearman's rank correlation.

**g**, Comparison of YF2 $IC_{50}$ values between germinal center (GC) and non-GC B-cell lymphoma cell lines. Data are presented as means with quartiles; comparisons were performed using an unpaired two-sided t-test ($n$ = 5–11).

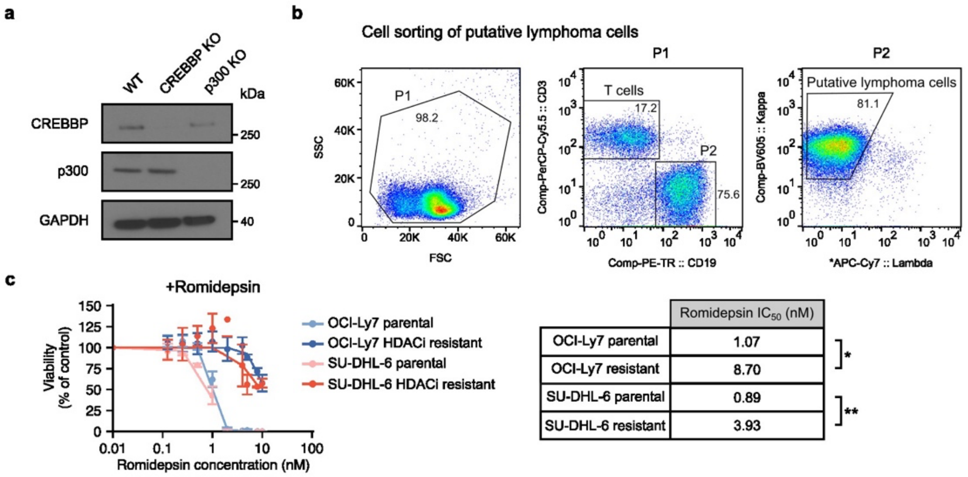


| | Romidepsin $IC_{50}$ (nM) | |
|---|---|---|
| OCI-Ly7 parental | 1.07 | * |
| OCI-Ly7 resistant | 8.70 | |
| SU-DHL-6 parental | 0.89 | ** |
| SU-DHL-6 resistant | 3.93 | |

**Supplementary Fig. S3: Generation and validation of experimental cell models.**

**a**, Immunoblot analysis of OCI-Ly7 wild-type and CREBBP/p300 knockout cells corresponding to **Fig. 1e**.

**b**, Representative gating strategy for isolation of primary follicular lymphoma (FL) cells corresponding to **Fig. 1f**.

**c**, Validation of histone deacetylase inhibitor (HDACi)-resistant cell lines. Parental and resistant lines were treated with increasing concentrations of romidepsin for six days, and viability was assessed. Data are presented as means ± s.e.m., normalized to vehicle controls. $IC_{50}$ values were compared using a Mann–Whitney U test ($n$ = 3). Asterisks (*) denote statistical significance: * = $P < 0.05$ and ** = $P < 0.01$.

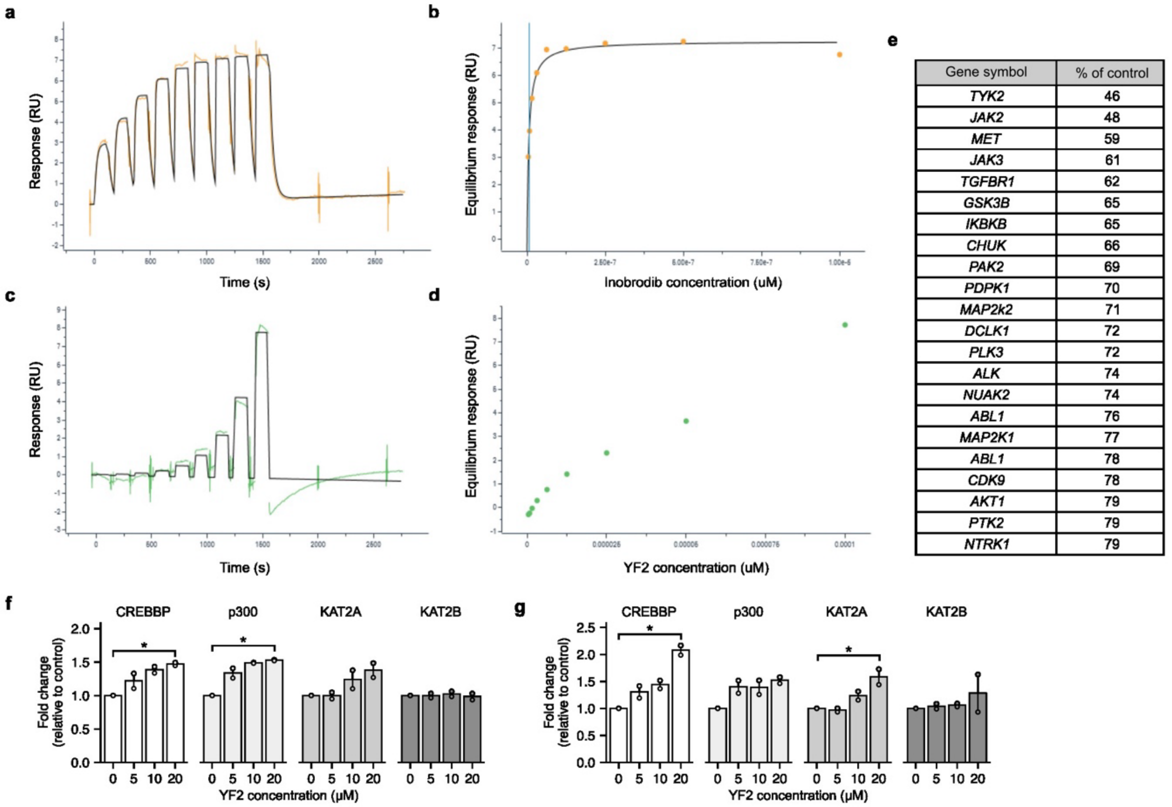


| Gene symbol | % of control |
|---|---|
| *TYK2* | 46 |
| *JAK2* | 48 |
| *MET* | 59 |
| *JAK3* | 61 |
| *TGFBR1* | 62 |
| *GSK3B* | 65 |
| *IKBKB* | 65 |
| *CHUK* | 66 |
| *PAK2* | 69 |
| *PDPK1* | 70 |
| *MAP2k2* | 71 |
| *DCLK1* | 72 |
| *PLK3* | 72 |
| *ALK* | 74 |
| *NUAK2* | 74 |
| *ABL1* | 76 |
| *MAP2K1* | 77 |
| *ABL1* | 78 |
| *CDK9* | 78 |
| *AKT1* | 79 |
| *PTK2* | 79 |
| *NTRK1* | 79 |

**Supplementary Fig. S4: YF2 bromodomain binding and selectivity profile.**
**a**, SPR sensorgram of inobrodib binding to the p300 bromodomain.
**b**, Steady-state affinity plot of inobrodib binding to the p300 bromodomain.
**c**, SPR sensorgram of YF2 binding to the p300 bromodomain.
**d**, Steady-state affinity plot of YF2 binding to the p300 bromodomain.
**e**, Kinase selectivity profiling of YF2. Data are presented as percent activity relative to vehicle controls; only kinases with <80% activity are shown ($n = 3$).
**f**, Cell-free HAT assay measuring global H3 acetylation across enzymes following 30 min YF2 exposure. Data are presented as fold change relative to controls (means ± s.e.m.); comparisons were performed using Kruskal–Wallis with Dunn's post-hoc test ($n = 2$).
**g**, Cell-free HAT assay measuring global H4 acetylation under the same conditions as **f** ($n = 2$).
Asterisks (*) denote statistical significance: * = $P < 0.05$.

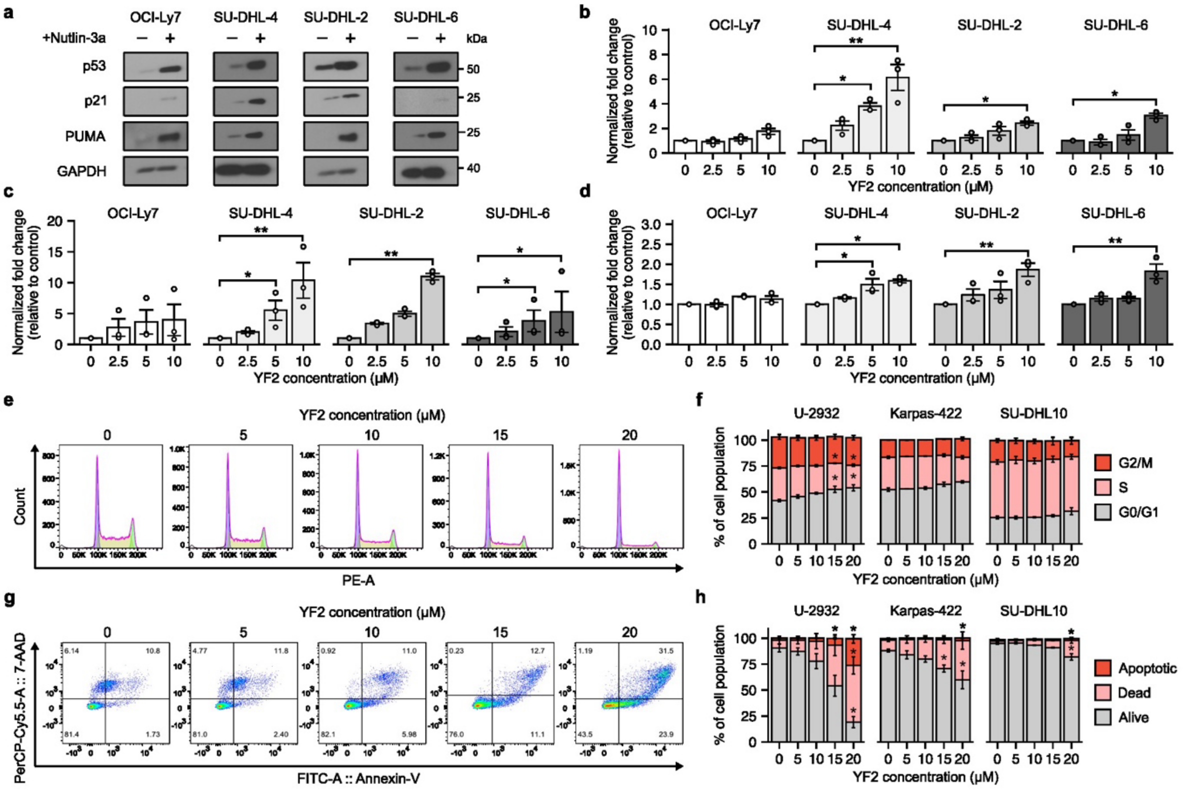


**Supplementary Fig. S5: Validation of p53 pathway activation and downstream phenotypes.**

**a**, Immunoblot analysis of p53 and downstream targets in B-cell lymphoma cell lines treated with the MDM2 inhibitor Nutlin-3a for 24 h ($n$ = 3).

**b**, Quantification of acetylated p53 levels shown in **Fig. 4d**. Data are normalized to total p53 and GAPDH and presented as fold change relative to controls (means ± s.e.m.). Statistical comparisons were performed using Kruskal–Wallis with Dunn's post-hoc test ($n$ = 3).

**c**, Quantification of p21 expression as described in **b** ($n$ = 3).

**d**, Quantification of PUMA expression as described in **b** ($n$ = 3).

**e**, Representative flow cytometry histograms from cell cycle analysis described in **Fig. 4g** ($n$ = 3).

**f**, Quantified cell cycle distribution following six-day YF2 treatment described in **Fig. 4g**. Data are presented as means ± s.e.m.; comparisons were performed using Kruskal–Wallis with Dunn's post-hoc test ($n$ = 3).

**h**, Quantified cell cycle distribution following six-day YF2 treatment described in **Fig. 4i**. Data are presented as means ± s.e.m.; comparisons were performed using Kruskal–Wallis with Dunn's post-hoc test ($n$ = 3).

Asterisks (*) denote statistical significance: * = $P < 0.05$ and ** = $P < 0.01$.

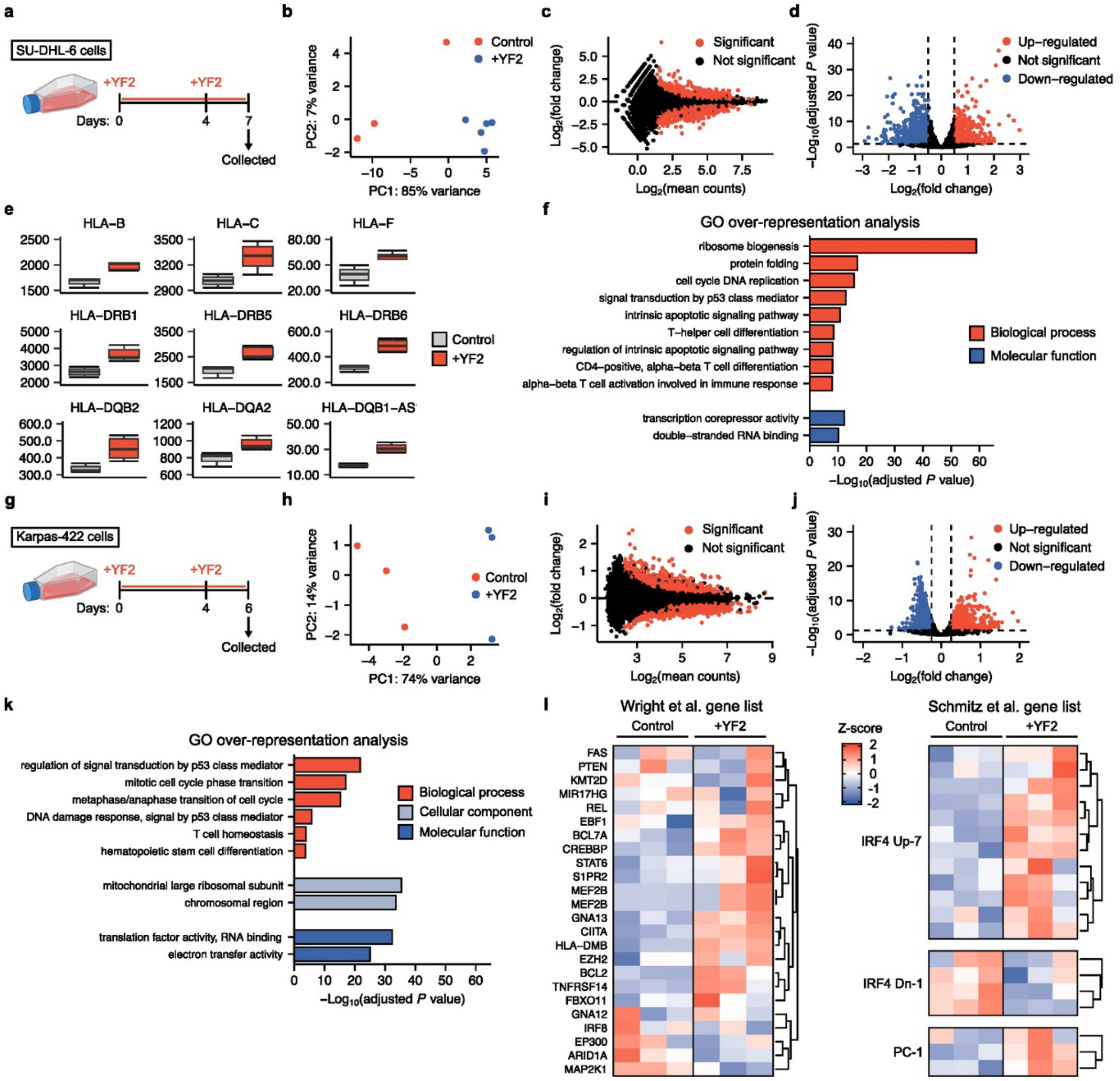


**Supplementary Fig. S6: YF2 induces transcriptional programs associated with immune regulation and lymphomagenesis.**

**a**, Schematic depicting bulk RNA sequencing (RNA-seq) workflow for *CREBBP*+*EP300*-mutated SU-DHL-6 cells treated with 10 μM YF2 for seven days before sample collection. Created with BioRender.com.

**b**, Principal component analysis of SU-DHL-6 RNA-seq samples demonstrating separation between YF2-treated and vehicle-treated groups ($n$ = 3–5).

**c**, MA plot showing global differential gene expression in SU-DHL-6 cells following YF2 treatment ($n$ = 3–5).

**d**, Volcano plot depicting significantly up- and down-regulated genes in YF2-treated SU-DHL-6 cells (adjusted $P < 0.05$; $n$ = 3–5).

**e**, Differential expression of antigen-presentation genes in SU-DHL-6 RNA-seq samples following YF2 treatment ($n$ = 3–5).

**f**, Gene ontology (GO) over-representation analysis of significantly altered genes in YF2-treated SU-DHL-6 cells. Select significant pathways (adjusted $P < 0.05$) are shown ($n = 3$–5).

**g**, Schematic depicting bulk RNA-seq workflow for *CREBBP+EP300*-mutated Karpas-422 cells treated with 8 μM YF2 for six days before sample collection. Created with BioRender.com.

**h**, Principal component analysis of Karpas-422 RNA-seq samples demonstrating separation between treatment groups ($n = 3$).

**i**, MA plot showing global differential gene expression in Karpas-422 cells following YF2 treatment ($n = 3$).

**j**, Volcano plot depicting significantly up- and down-regulated genes in YF2-treated Karpas-422 cells (adjusted $P < 0.05$; $n = 3$).

**k**, GO over-representation analysis of significantly altered genes in YF2-treated Karpas-422 cells. Select significant pathways (adjusted $P < 0.05$) are shown ($n = 3$).

**l**, Heatmap of Karpas-422 RNA-seq samples showing YF2-induced differential expression of genes associated with the EZB DLBCL subtype defined by Wright et al. ($n = 3$).

**m**, Heatmap of Karpas-422 RNA-seq samples showing YF2-induced differential expression of EZB-associated gene signatures defined by Schmitz et al. ($n = 3$).

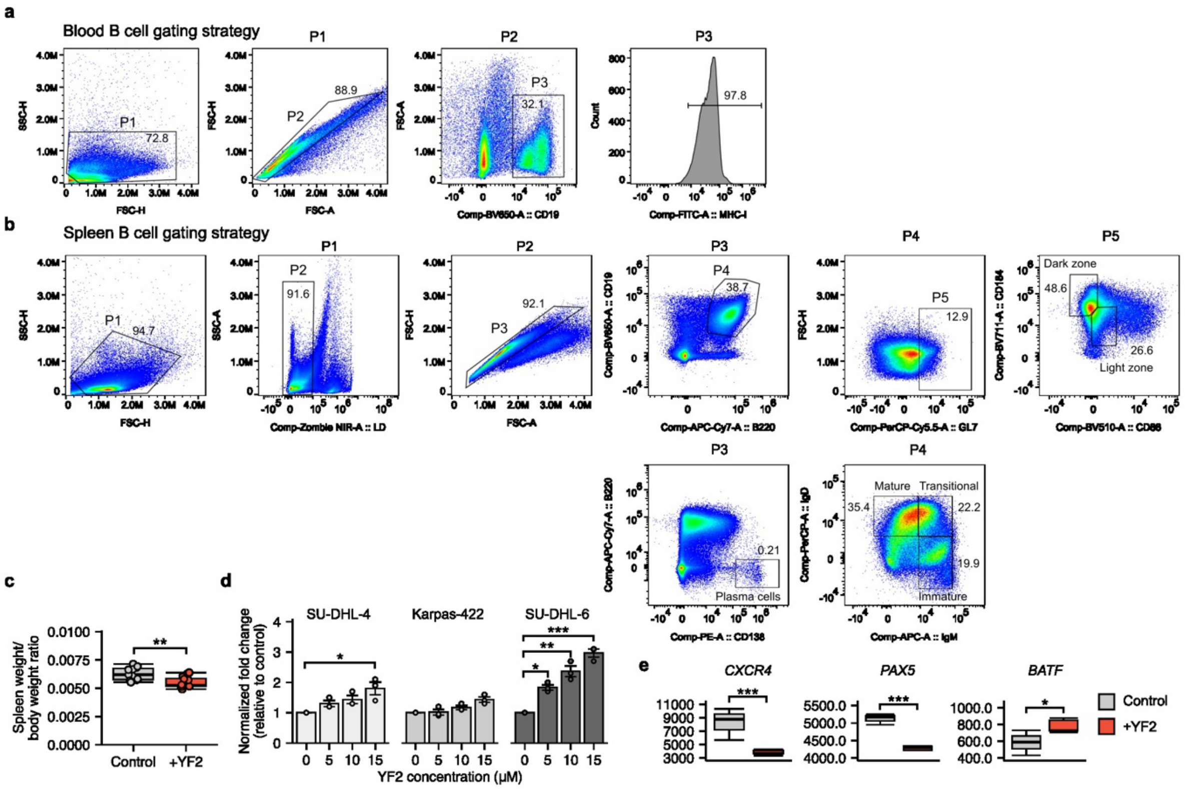


**Supplementary Fig. S7: Additional analyses of YF2-induced B-cell differentiation and germinal center remodeling.**

**a**, Representative flow cytometry gating strategy for peripheral blood B cells from the non-engrafted BALB/c mouse experiment.

**b**, Representative flow cytometry gating strategy for splenic B cells from the non-engrafted BALB/c mouse experiment.

**c**, Spleen-to-body weight ratios in vehicle- and YF2-treated mice. Data are presented as means ± s.e.m.; groups were compared using an unpaired two-sided t-test (*n* = 10–11).

**d**, Quantification of PRDM1 protein expression shown in **Fig. 5l**. Data are presented as normalized fold change relative to vehicle controls (means ± s.e.m.). Statistical comparisons were performed using Kruskal–Wallis with Dunn's post-hoc test (*n* = 3).

**e**, Differential expression of germinal center regulatory genes in SU-DHL-6 RNA-seq samples following YF2 treatment (*n* = 3–5).

Asterisks (*) denote statistical significance: * = $P < 0.05$, ** = $P < 0.01$, and *** = $P < 0.001$.

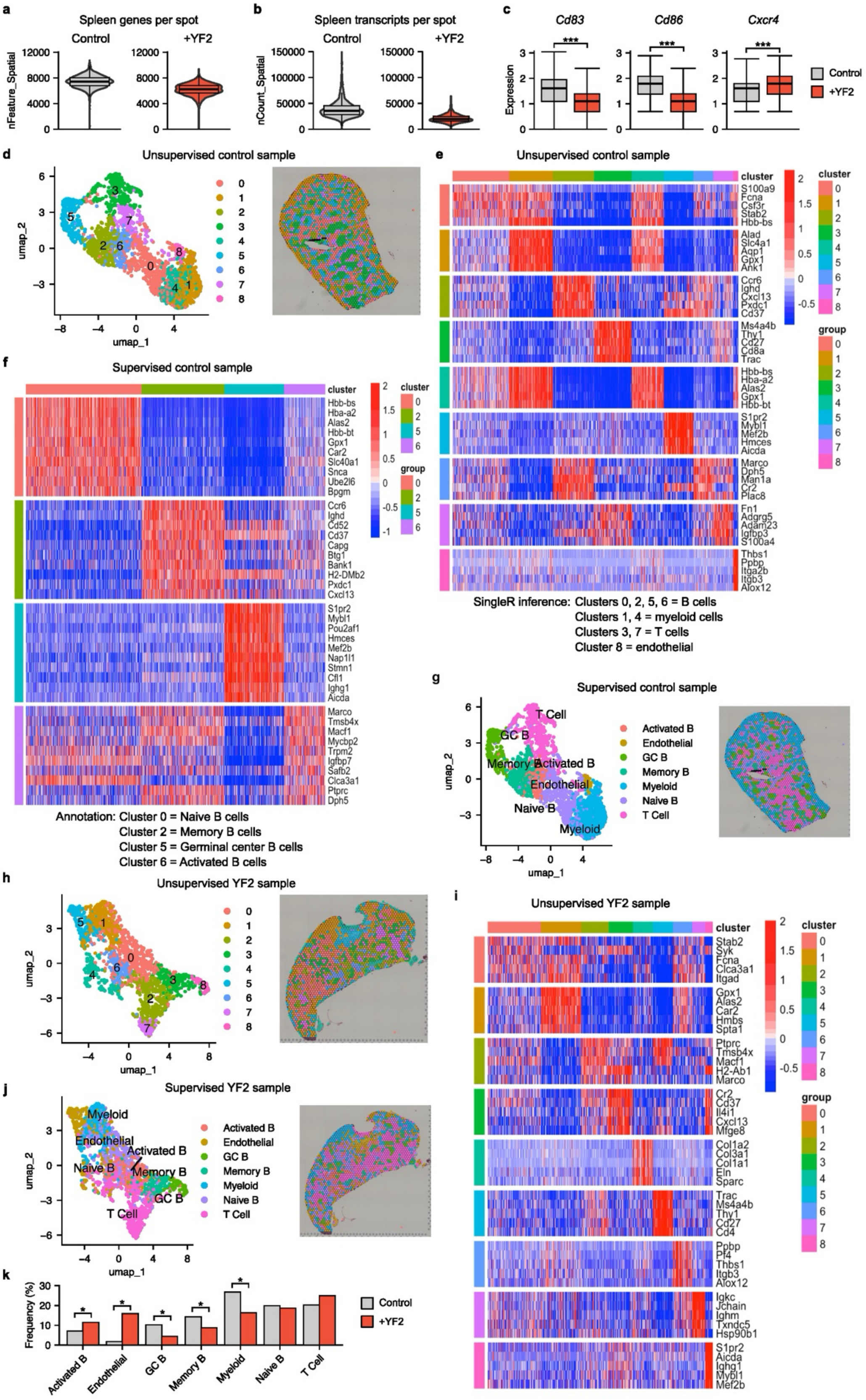
a
Spleen genes per spot
Control
+YF2
nFeature_Spatial
b
Spleen transcripts per spot
Control
+YF2
nCount_Spatial
c
Cd83
Cd86
Cxcr4
***
Expression
Control
+YF2
d
Unsupervised control sample
umap_1
umap_2
e
Unsupervised control sample
cluster
group
S100a9
Fcna
Csf3r
Stab2
Hbb-bs
Alad
Slc4a1
Aqp1
Gpx1
Ank1
Ccr6
Ighd
Cxcl13
Pxdc1
Cd37
Ms4a4b
Thy1
Cd27
Cd8a
Trac
Hbb-bs
Hba-a2
Alas2
Gpx1
Hbb-bt
S1pr2
Mybl1
Mef2b
Hmces
Aicda
Marco
Dph5
Man1a
Cr2
Plac8
Fn1
Adgrg5
Adam23
Igfbp3
S100a4
Thbs1
Ppbp
Itga2b
Itgb3
Alox12
SingleR inference: Clusters 0, 2, 5, 6 = B cells
Clusters 1, 4 = myeloid cells
Clusters 3, 7 = T cells
Cluster 8 = endothelial
f
Supervised control sample
cluster
group
Hbb-bs
Hba-a2
Alas2
Hbb-bt
Gpx1
Car2
Slc40a1
Snca
Ube2l6
Bpgm
Ccr6
Ighd
Cd52
Cd37
Capg
Btg1
Bank1
H2-DMb2
Pxdc1
Cxcl13
S1pr2
Mybl1
Pou2af1
Hmces
Mef2b
Nap1l1
Stmn1
Cfl1
Ighg1
Aicda
Marco
Tmsb4x
Macf1
Mycbp2
Trpm2
Igfbp7
Safb2
Clca3a1
Ptprc
Dph5
Annotation: Cluster 0 = Naive B cells
Cluster 2 = Memory B cells
Cluster 5 = Germinal center B cells
Cluster 6 = Activated B cells
g
Supervised control sample
T Cell
GC B
Memory B
Activated B
Endothelial
Naive B
Myeloid
Activated B
Endothelial
GC B
Memory B
Myeloid
Naive B
T Cell
umap_1
umap_2
h
Unsupervised YF2 sample
umap_1
umap_2
i
Unsupervised YF2 sample
cluster
group
Stab2
Syk
Fcna
Clca3a1
Itgad
Gpx1
Alas2
Car2
Hmbs
Spta1
Ptprc
Tmsb4x
Macf1
H2-Ab1
Marco
Cr2
Cd37
Il4i1
Cxcl13
Mfge8
Col1a2
Col3a1
Col1a1
Eln
Sparc
Trac
Ms4a4b
Thy1
Cd27
Cd4
Ppbp
Pf4
Thbs1
Itgb3
Alox12
Igkc
Jchain
Ighm
Txndc5
Hsp90b1
S1pr2
Aicda
Ighg1
Mybl1
Mef2b
j
Supervised YF2 sample
Myeloid
Endothelial
Activated B
Naive B
Memory B
GC B
T Cell
Activated B
Endothelial
GC B
Memory B
Myeloid
Naive B
T Cell
umap_1
umap_2
k
Frequency (%)
Control
+YF2
Activated B
Endothelial
GC B
Memory B
Myeloid
Naive B
T Cell

**Supplementary Fig. S8: Spatial transcriptomic profiling of YF2-treated murine spleens.**

**a**, Gene counts per spatial transcriptomic spot in vehicle- and YF2-treated murine spleens (*n* = 1).

**b**, Transcript counts per spatial transcriptomic spot in vehicle- and YF2-treated murine spleens (*n* = 1).

**c**, Quantification of spatially differential gene expression across treatment groups. Spatial feature expression was normalized across samples for each target (*n* = 1).

**d**, Dimensionality-reduced and spatially resolved plots of the vehicle-treated spleen following unsupervised clustering.

**e**, Heatmap showing the top five differentially expressed genes per cluster after unsupervised clustering of the vehicle-treated spleen. Clusters were annotated as B cells (0, 2, 5, and 6), myeloid cells (1 and 4), T cells (3 and 7), and endothelial cells (8).

**f**, Heatmap showing the top 10 differentially expressed genes per cluster following supervised clustering of the vehicle-treated spleen. B-cell clusters were further annotated as naïve (cluster 0), memory (cluster 2), germinal center (cluster 5), and activated B cells (cluster 6).

**g**, Dimensionality-reduced and spatially resolved plots of the vehicle-treated spleen following supervised clustering and annotation.

**h**, Dimensionality-reduced and spatially resolved plots of the YF2-treated spleen following unsupervised clustering.

**i**, Heatmap showing the top five differentially expressed genes per cluster after unsupervised clustering of the YF2-treated spleen.

**j**, Dimensionality-reduced and spatially resolved plots of the YF2-treated spleen following supervised clustering and annotation.

**k**, Differential spatial abundance of splenic cell populations following treatment. Cell-type abundance comparisons were assessed using Fisher's exact test with Benjamini–Hochberg correction.

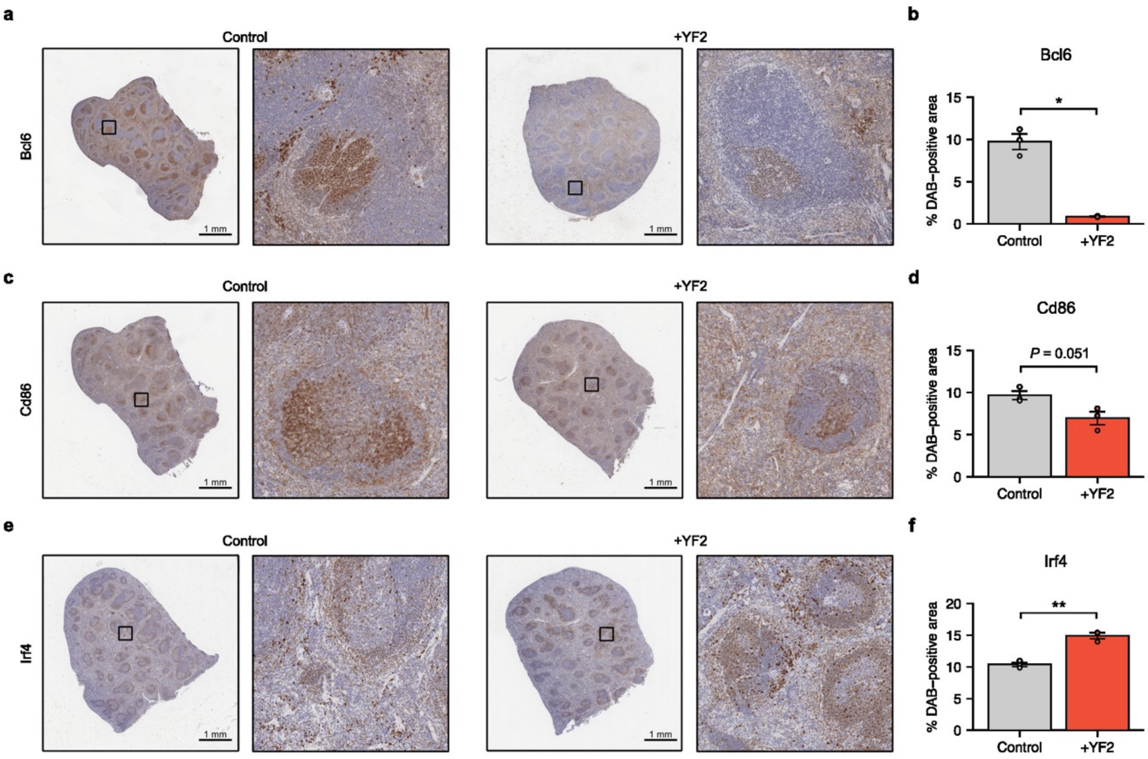


**Supplementary Fig. S9: YF2 alters germinal center organization and B-cell differentiation in murine spleens.**

**a**, Representative immunohistochemical staining of Bcl6 in murine spleens following *in vivo* YF2 treatment ($n$ = 3).

**b**, Quantification of Bcl6-positive DAB staining, measured as percent positive area across five fields per spleen ($n$ = 3).

**c**, Representative immunohistochemical staining of Cd86 in murine spleens following *in vivo* YF2 treatment ($n$ = 3).

**d**, Quantification of Cd86-positive DAB staining, measured as percent positive area across five fields per spleen ($n$ = 3).

**e**, Representative immunohistochemical staining of Irf4 in murine spleens following *in vivo* YF2 treatment ($n$ = 3).

**f**, Quantification of Irf4-positive DAB staining, measured as percent positive area across five fields per spleen ($n$ = 3).

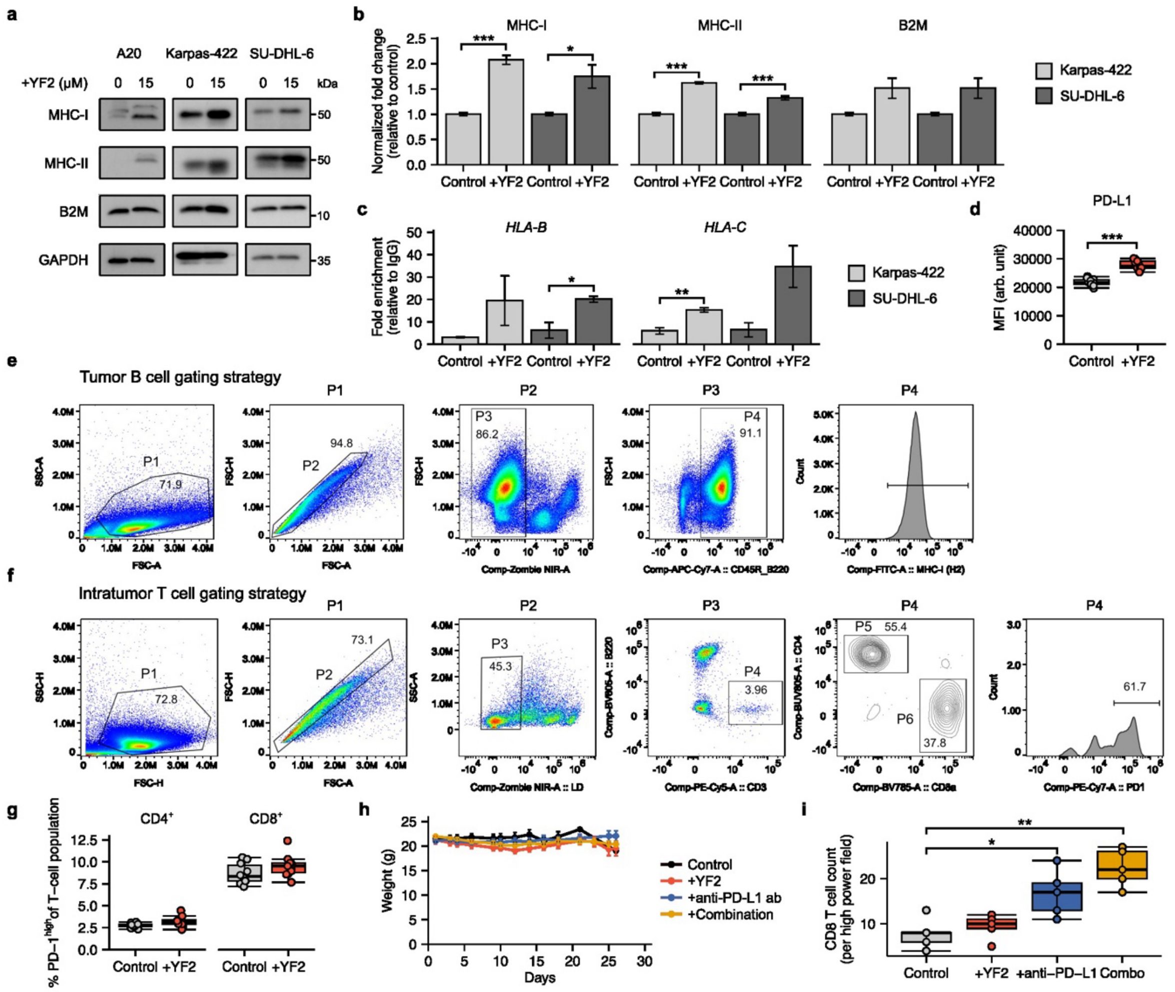


**Supplementary Fig. S10: Additional analyses of antigen presentation and immunotherapy response following YF2 treatment.**

**a**, Immunoblot analysis of major histocompatibility complex class I (MHC-I), MHC-II, and beta-2-microglobulin (B2M) expression in B-cell lymphoma cell lines treated with YF2 (*n* = 3).

**b**, Quantification of MHC-I, MHC-II, and B2M protein expression shown in **a**. Data are presented as normalized fold change relative to vehicle controls (means ± s.e.m.). Statistical comparisons were performed using unpaired two-sided t-tests (*n* = 3).

**c**, H3K27ac chromatin immunoprecipitation showing enrichment of *HLA-B* and *HLA-C* expression in SU-DHL-6 cells treated with 10 μM YF2 for 24 h. Data are presented as fold enrichment relative to IgG controls (means ± s.e.m.); comparisons were performed using unpaired two-sided t-tests (*n* = 3).

**d**, Cell-surface PD-L1 expression on peripheral blood B cells following YF2 treatment in the A20 lymphoma mouse model, assessed by flow cytometry. Data are presented as mean fluorescence intensity (MFI; means ± s.e.m.); comparisons were performed using unpaired two-sided t-tests (*n* = 9–10).

**e**, Representative flow cytometry gating strategy for tumor-associated B cells from the A20 lymphoma mouse experiment.

**f**, Representative flow cytometry gating strategy for tumor-associated T cells from the A20 lymphoma mouse experiment.

**g**, Frequency of PD-1$^{hi}$ T cells following YF2 treatment in the A20 lymphoma mouse experiment. Data are presented as percentages of $CD4^+$ or $CD8^+$ T-cell populations (means ± s.e.m.); comparisons were performed using unpaired two-sided t-tests ($n$ = 10–11).

**h**, Body weight changes during combination treatment in the A20 lymphoma mouse experiment. Data are presented as means ± s.e.m.; treatment groups were compared using linear regression analysis ($n$ = 9–10).

**i**, Quantification of intratumoral $CD8^+$ cells per high-power field shown in **Fig. 6l**. Data are presented as means with quartiles; statistical comparisons were performed using one-way ANOVA with Tukey's HSD post-hoc test ($n$ = 3).

Asterisks (*) denote statistical significance: * = $P < 0.05$, ** = $P < 0.01$, and *** = $P < 0.001$.

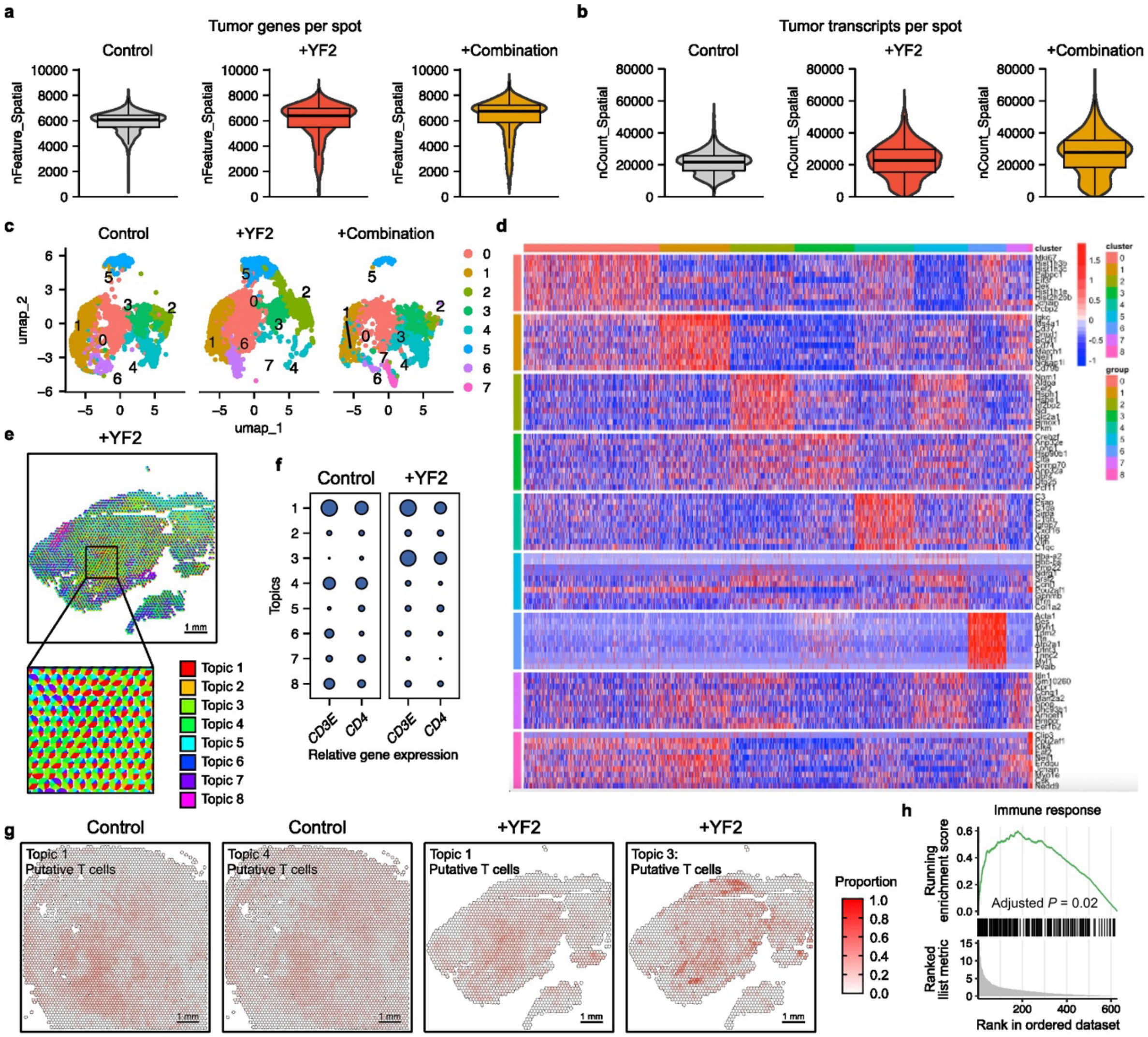


**Supplementary Fig. S11: Spatial transcriptomic profiling of YF2-treated murine tumors.**

**a**, Gene counts per spatial transcriptomic spot across murine tumor treatment groups (*n* = 1).

**b**, Transcript counts per spatial transcriptomic spot across murine tumor treatment groups (*n* = 1).

**c**, Dimensionality reduction of integrated tumor samples across treatment groups.

**d**, Heatmap showing the top 10 differentially expressed genes per cluster following supervised clustering of integrated tumor samples.

**e**, Spatial organization of inferred cell populations (“topics”) identified by latent Dirichlet allocation modeling of tumor samples (*n* = 1).

**f**, Relative expression of T-cell marker genes across inferred spatial topics in vehicle- and YF2-treated tumors. Topics 1 and 4 were annotated as putative T cells in control tumors, whereas topics 1 and 3 were annotated as putative T cells in YF2-treated tumors (*n* = 1).

**g**, Spatial distribution of putative T-cell populations in vehicle- and YF2-treated tumors (*n* = 1).

**h**, Gene set enrichment analysis (GSEA) of immune response pathways using differentially expressed genes from topic 3 in the YF2-treated tumor sample (*n* = 1).
